# Cryo-EM Reveals the Mechanochemical Cycle of Reactive Full-length Human Dynein-1

**DOI:** 10.1101/2024.05.01.592044

**Authors:** Pengxin Chai, Jun Yang, Indigo C. Geohring, Steven M. Markus, Yue Wang, Kai Zhang

## Abstract

Dynein-driven cargo transport plays pivotal roles in diverse cellular activities, central to which is dynein’s mechanochemical cycle. Here, we performed a systematic cryo-electron microscopic investigation of the conformational landscape of full-length human dynein-1 in reaction, under various nucleotide conditions, on and off microtubules. Our approach reveals over 40 high-resolution structures, categorized into eight states, providing a dynamic and comprehensive view of dynein throughout its mechanochemical cycle. The novel intermediate states reveal important mechanistic insights into dynein function, including a ‘backdoor’ phosphate release model that coordinates linker straightening, how microtubule binding enhances ATPase activity through a two-way communication mechanism, and the crosstalk mechanism between AAA1 and the regulatory AAA3 site. Our findings also lead to a substantially revised model for the force-generating powerstroke and reveal a means by which dynein exhibits unidirectional stepping. These results substantially improve our understanding of dynein and provide a more complete model of its mechanochemical cycle.

## Introduction

Dyneins are large cytoskeletal motor complexes that move toward the minus ends of microtubules (MTs)(1). Cytoplasmic dynein-1 is the sole cytoplasmic isoform that drives retrograde intracellular cargo transport(2), organelle positioning(3), and cellular morphogenesis(4), while dynein-2 and axonemal dyneins mediate intraflagellar transport(5) and ciliary motility(6). Unlike the kinesin family which encompasses 45 different types of predominantly plus end-directed motors in humans(7), a single cytoplasmic dynein-1 (dynein) is responsible for transporting diverse cellular cargos including membrane-bound organelles, RNAs, proteins, and viruses(2). This is enabled by the processive retrograde transport machinery that includes dynein, dynactin, and a cargo adaptor (DDA complex)(8–15), and is further regulated by cofactors such as LIS1(16–23) and Ndel1(24–27). Owing to its fundamental roles in a variety of cellular activities, mutations in dynein and its cofactors have been implicated in many human diseases, including neurodegenerative(28) and neurodevelopmental disorders(29–31), and viral infections(32).

The dynein complex comprises two copies of the heavy chain (DHC) and several accessory chains. Each DHC consists of a tail domain, responsible for binding dynactin, adaptors and accessory chains, and a motor domain that is essential for motility(2, 33–35). The motor domain includes an AAA+ (ATPase associated with cellular activities) ring that consists of six non-identical AAA+ domains, as well as a linker, coiled-coil stalk, and buttress appendages, a microtubule-binding domain (MTBD) located at the tip of the stalk, and a C-terminal domain (CTD). The motor domain possesses ATPase activity and is responsible for converting the chemical energy stored in ATP into mechanical processes, collectively termed the mechanochemical cycle(35).

The mechanochemical cycle of dynein is a fundamental process that underlies all dynein-mediated cellular activities. Our current understanding of this cycle primarily stems from biochemical(36–39), biophysical(40–43), and structural studies(35, 44–49) conducted on dynein motor domain fragments. Within the motor domain, AAA1 serves as the primary site for ATP hydrolysis(36, 38). Throughout the catalysis cycle, the AAA+ ring adopts alternating closed and open conformations that are critical for force generation and microtubule-based motility(35, 50). The linker domain can change from a bent to a straight conformation in a nucleotide-dependent manner, which has been deemed responsible for the force-generating “powerstroke”(37, 42, 49, 51, 52). When straight, the linker docks onto a conserved site of AAA5(48, 53). However, a recent study of axonemal outer-arm dynein revealed that the linker can dock at a distinct site close to AAA4, termed “post-2” (compared to the classical “post-1” docking mode(54)). Whether dynein-1 also samples the post-2 state, and the role of post-2 during dynein’s mechanochemical cycle is unclear. Both kinetic and structural evidence indicate that the release of phosphate (Pi) that is coupled with the opening of the AAA+ ring, and the AAA1 pocket specifically, triggers the straightening of the linker(35, 37, 42, 50). However, the mechanism by which Pi is released from AAA1 remains elusive. Does it follow a “molecular backdoor” mechanism akin to myosin(55) and actin(56) which only requires minimal local conformational change, or does it require complete pocket opening? As a result, the correlation between Pi release, pocket opening, and linker straightening remains poorly understood(35).

In contrast to the other two cytoskeletal motor proteins, kinesin and myosin, which bind to their respective tracks with their ATP hydrolysis domains(55, 57), the MTBD of dynein is 24 nm away from the AAA1 domain(33). The nucleotide-bound states and pocket conformations of AAA1 have been shown to regulate the MTBD affinity for MTs(47–49). Meanwhile, the MTs provide a track for the unidirectional stepping of dynein(40, 43, 58) and stimulate the AAA1 ATPase activity(47, 59). Communication between AAA1 and the MTBD is mediated by the buttress(46, 47) and the 15-nm-long stalk coiled-coil (stalk-CC)(60–62). The coupled conformational change of the buttress-stalk can leads to multiple registries within the stalk-CC that correspond to different MT-binding affinities(41, 63–67), which is regulated by both AAA1 pocket conformations(35) and MT-binding(62, 68, 69). However, due to the limited structural information available for dynein on and off MTs, the precise mechanism of communication between dynein and MTs via the buttress-stalk remains unclear. Therefore, how dynein’s mechanochemical cycle is accelerated by MTs(70), and how dynein achieves unidirectional motility along MTs during the mechanochemical cycle(58) requires further investigation.

Previous studies have elucidated the structures of truncated dynein(44, 46–49), full-length “phi” dynein-1(15) (similar to the Greek letter “*ϕ*”(71)), phi dynein-2(72), outer-arm dynein (OAD)-MT(54), and the DDA-MT(8, 22) complex, each captured under specific nucleotide conditions. Due to either limited resolution, or the lack of structural dynamics inherent to these conventional approaches (e.g., induced by non-hydrolyzable ATP), these structures provide valuable but limited insight into the atomic-level details of the mechanochemical cycle of full-length dynein. Here, we employed strategies to capture snapshots of the complete conformational landscape of full-length dynein progressing through its natural mechanochemical cycle. First, we imaged full-length dynein in the presence of ATP (to visualize dynein “in reaction”) as well as other nucleotide conditions, which allowed us to obtain structures of phi dynein and novel intermediate states from open/uninhibited dynein with much-improved resolution (2.2-3.6 Å). Furthermore, we imaged reconstituted MT-bound dynein complexes in different nucleotide conditions and obtained structures of the motor domains at high resolution (2.9-3.6 Å). This collection of motor domain structures (over 40) provides critical insight into several longstanding questions in the field. Our results reveal a “backdoor” channel through which Pi is released from AAA1, a mechanism to account for linker straightening, and how communication between AAA1 and the MTBD occurs, which we support with mutagenesis and functional studies. Comparison of dynein motor domains on and off MTs reveals the structural basis for enhanced ATPase activity upon MT-binding, the role of linker-AAA+ ring interactions in promoting unidirectional stepping, and a revised model for the dynein powerstroke. Our large-scale structural survey also reveals a crosstalk mechanism between AAA1 and the regulatory AAA3 site. In summary, our comprehensive structural analysis of full-length dynein coupled with functional investigations significantly advances our understanding of the dynein mechanochemical cycle.

## Results

### Structural determination of full-length human dynein-1 off and on MTs

To explore the conformational landscape of dynein throughout its mechanochemical cycle (“in reaction”), we purified full-length human dynein-1 from insect cells (which had a roughly equal mixture of phi and open states) and prepared cryo-EM grids in the presence of 5 mM ATP-Mg^2+^ (**Fig. 1a**). We collected a large cryo-EM dataset and performed extensive in-silico 3D classification analysis followed by high-resolution refinement (**Extended Data Fig. 1, 2 and Supplementary Table 1**). We obtained high-resolution cryo-EM maps of full-length phi dynein (**Fig. 1c, Supplementary Video 1**), with the motor domain at 2.2 Å average resolution, and the majority of the tail region better than 4 Å. This significantly improved resolution compared to the previous one(15) allowed us to build a full atomic model for phi dynein. In addition to the phi dynein motor, we observed 10 distinct conformational states of the open motor domain, revealing snapshots of reactive intermediates during dynein’s mechanochemical cycle (**Fig. 1d**). The high-resolution structure of the motor domain provides unprecedented details, including holes within aromatic side chains, water molecules, and ions interacting with the nucleotide (**Fig. 1f**).

**Fig. 1.**
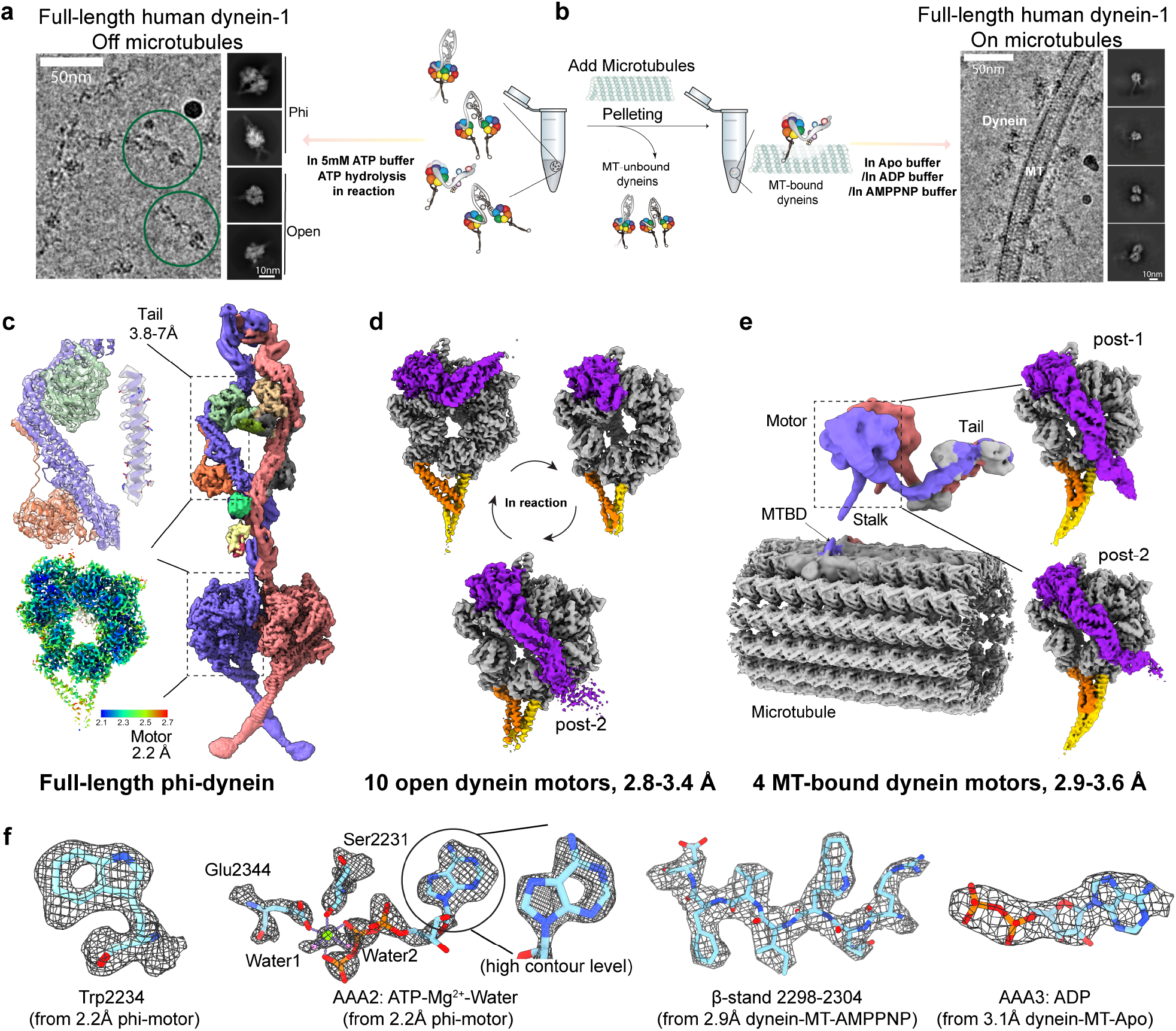
Cryo-EM analysis of full-length human dynein-1 off and on microtubules. **(a,b)** Schematic of cryo-EM pipeline for determination of dynein structures in reaction with ATP without MTs (a), or with MTs in different nucleotide conditions (b). **(c)** Cryo-EM reconstruction of phi dynein with the 3.8-7.0 Å tail region and 2.2 Å motor domain. **(d)** Representative cryo-EM reconstructions of the motor domain in different states from open dynein. The motor ring, linker, buttress, and stalk are colored grey, purple, orange, and yellow, respectively. **(e)** Cryo-EM map of full-length two-headed dynein bound to MTs in ADP conditions at 10 Å resolution, with locally refined motor domains at 2.9-3.6 Å resolution. **(f)** High-resolution features are shown by representative density of amino acid residues and ligands.

High-resolution structural determination of MT-bound dynein has been a challenge in the field. To address this, we recently developed a micrograph-based MT-signal subtraction method(73, 74), leading to the successful reconstruction of OAD-MT and DDA-MT structures(8, 22, 54). To capture complete MT-bound dynein-1 structures, we reconstituted full-length dynein-1 on microtubules in apo, ADP, and AMPPNP conditions, all of which increase dynein affinity for MTs(75) (**Fig. 1b, Extended Data Fig. 3 and Supplementary Table 2**). We obtained a structure of two-headed full-length dynein bound to MTs at a resolution of 10 Å (**Fig. 1e**). Focused classification and refinement of the individual motor domains resulted in 4 distinct MT-bound motor domain structures at 2.9-3.6 Å (**Fig. 1e, and Supplementary Video 2**).

### Overview of the conformational landscape of microtubule-unbound dynein

By focusing on the following four structural features, we obtained 10 distinct motor domain structures from phi and open dynein: (1) the pocket conformations of AAA1, (2) and of AAA3, (3) the linker position, and (4) the buttress-stalk conformations. By primarily focusing on AAA1 and the overall AAA+ ring arrangement, these 10 structures were further categorized into 8 major states (state-1 to state-8) that represent distinct points of the mechanochemical cycle (**Fig. 2a**). The sequence of the different states was determined by AAA1 pocket dynamics and apparent nucleotide-bound states, as well as previous models of the dynein cycle(35), despite a lack of time-resolution in our cryo-EM analysis. Among the 8 states, 4 of them are novel: state-3, -4, -5, - Below we describe in detail the salient features of each state, which are also summarized in **Table 1**.

**Table 1.** Features of the motor domains in different states.

| Description | Buffer | State | AAA1 & motor ring | Linker | Buttress | Stalk-CC | MTBD affinity with MT | Analogous structures from published PDBs |
| --- | --- | --- | --- | --- | --- | --- | --- | --- |
| Full length human dynein-1 unbound from MTs | 5mM ATP | State-0 | open | straight, docked | straight | bulging | low | N/A |
|  | 5mM ATP | State-1 Phi-motor | closed | bent, docked | straight | bulging | low | 5nug, 6rla |
|  | 5mM ATP | State-2 | closed | bent, docked | straight | bulging | low | 4rh7 |
|  | 5mM ATP | State-3 | closed | bent, undocked | straight | bulging | low | N/A |
|  | 5mM ATP | State-4 | closed | straight, undocked | straight | bulging | low | N/A |
|  | 5mM ATP | State-5 | intermediate open | straight, undocked | intermediate | intermediate | intermediate | N/A |
|  | 5mM ATP | State-6 | intermediate open | straight, undocked | intermediate | intermediate | intermediate | N/A |
|  | 5mM ATP | State-7 | open | straight, docked at post-2 | kink | canonical | high | 3vkg*, 4w8f, 8fcy |
|  | apo | State-8 | wide open | straight, docked at post-1 | kink | canonical | high | 4aki |
| Full length human dynein-1 bound to MT | 5mM ADP | State-7 | open | straight, docked at post-1 | kink | canonical | high | N/A |
|  | 5mM ADP | State-7 | open | straight, docked at post-2 | kink | canonical | high | N/A |
|  | 5mM AMPPNP | State-7 | open | straight, docked at post-1 | kink | canonical | high | 7z8g |
|  | apo | State-8 | wide open | straight, docked at post-1 | kink | canonical | high | 7k58, 7k5b |

**Fig. 2.**
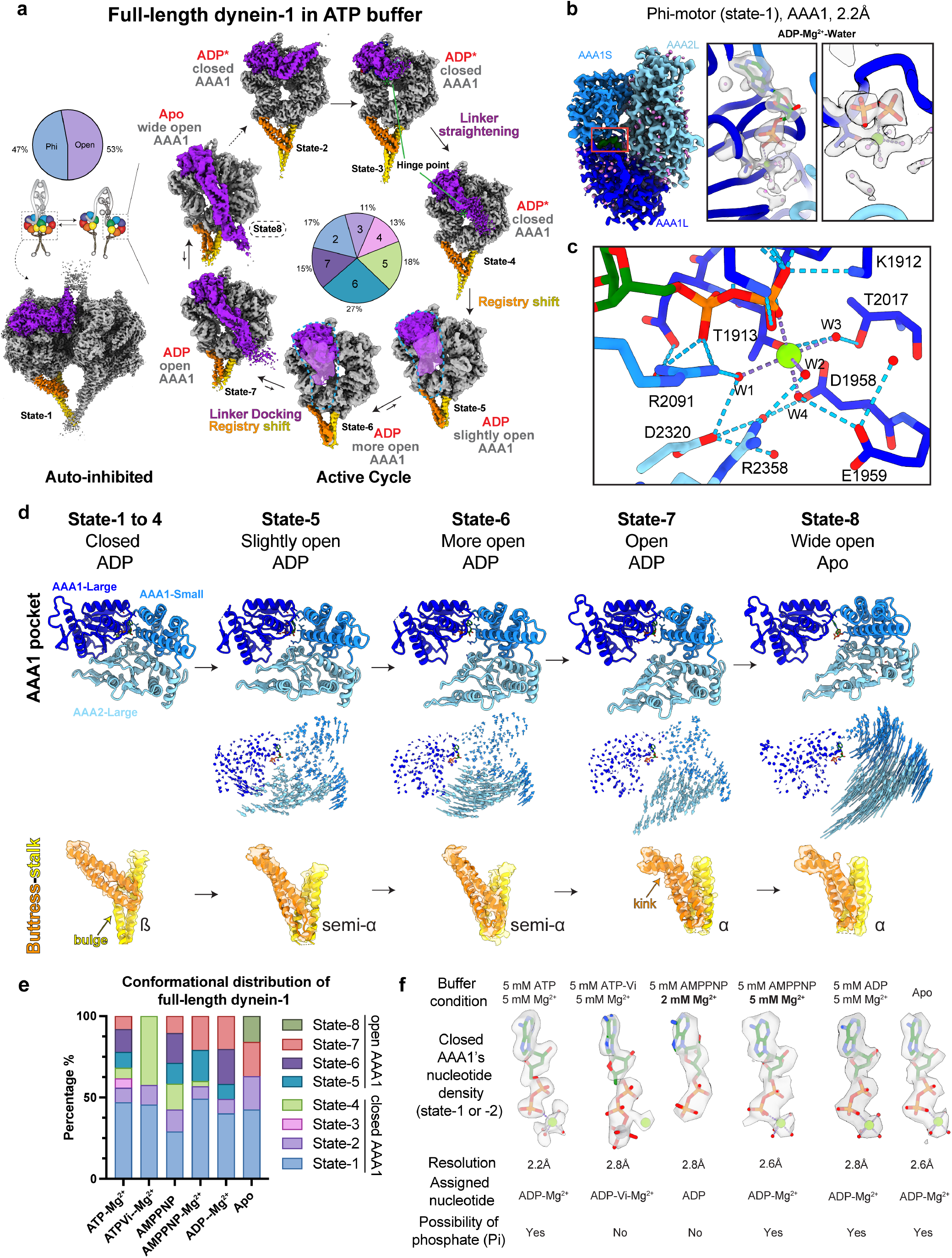
The conformational landscape of MT-unbound dynein motor domains in reaction. **(a)** Eight major states of the dynein motor domain during the catalytic cycle are divided into auto-inhibited states (left) and states in the active cycle (right). Arrows indicate the proposed catalytic pathway supported by the literature as well as apparent AAA+ ring conformations. Dashed arrows show the possibility of reverse pathway. Percentages indicate the proportion of particles in each state. The apo AAA1 pocket state (state-8) is present at a very low proportion in the presence of ATP but is highly represented in apo conditions (shown is the structure obtained in apo condition). The linkers from states-5 and -6 are low passed filtered to 12Å and shown at low contours. Only one representative sub-class in each state is shown. The color scheme is consistent with Fig. 1d. **(b)** Cryo-EM map of AAA1 pocket from 2.2 Å phi motors. AAA1L, AAA1S, and AAA2L are colored blue, light blue, and sky blue, respectively. On the right, carved transparent grey density is displayed for ADP (green), Mg^2+^ (light green), and water molecules (violet). **(c)**Side chain residues involved in hydrogen-bonding networks (dashed lines) at the AAA1 active site are displayed and colored by heteroatom. **(d)** Cartoon models showing the AAA1 dynamics and the coupled buttress-stalk conformational states. The isolated AAA1 pockets from different states are aligned on AAA1L. Vector maps depicting pairwise alpha carbon interatomic distances between the AAA1 domains of adjacent states depict the three-dimensional trajectories during the pocket opening process. The buttress-stalk regions are highlighted to show how the stalk-CC registries are coupled to motor ring states. **(e)** Plot depicting proportions of the various motor domain conformations from full-length dynein-1 in different nucleotide conditions. **(f)** EM densities for nucleotides bound to the closed AAA1 pockets under indicated nucleotide conditions.

#### States-1, -2, -3, and -4

These motor domains all have a closed AAA1 pocket and are in low-MT binding affinity states, as determined by the buttress and stalk conformations(35). State-1 (phi-motor) has the highest resolution (2.2 Å), enabling accurate modeling and refinement of ADP, magnesium, and water molecules at the active site (**Fig. 2b, c**). States-2, -3, and -4, resolved at 2.8 Å, 3.0 Å, and 2.8 Å, respectively, all have clear ADP-Mg^2+^ density in the AAA1 pocket (**Extended Data Fig. 2**). However, since these structures were solved in the presence of ATP, we cannot rule out the possibility of a traveling Pi (weak densities that cannot be well resolved) inside the pocket. Among all these states, AAA3 is in the open conformation with ADP bound. Unlike the autoinhibited state-1, states-2, -3, and -4 are actively engaged in the mechanochemical cycle and have different linker conformations. Specifically, state-2 possesses a bent linker that is stably docked between AAA2 and 3, similar to state-1 and previously reported pre-powerstroke motor structures (PDB: 4rh7(49), 5nug(15), 6rla(72), 8pqy(22)). State-4 has a straight “hinge helix”, which connects the N- and C-terminal regions of the linker, and an overall straight linker; however, the linker does not appear to be stably docked (apparent from poor electron density for N-terminal region), likely due to the closed AAA+ ring configuration in which the docking site at AAA5 is not exposed. The straight and undocked linker, and the apparent low MT binding affinity for state-4 suggest that this state is an intermediate during the dynein powerstroke. Finally, the hinge helix in state-3 adopts a bent conformation, while the rest of the linker region is flexible, indicating that state-3 represents an averaged map of many transient conformations between state-2 and state-4.

#### States-5, -6, and -7

All of these dyneins have ADP-bound to open AAA1 pockets, and account for about 60% of the motor states during the active ATP cycle, indicating that ADP release from AAA1 pocket is rate-limiting. All three of these states exhibit varying degrees of pocket opening that are correlated with different buttress-stalk conformations (**Fig. 2d**), consistent with the model whereby AAA1 pocket dynamics regulate MT affinity via the stalk-CC(35, 47–49). The AAA3 pocket for these states is primarily in an ATP-bound state with a closed conformation (**Extended Data Fig. 2**), indicating that AAA3 only undergoes nucleotide exchange when AAA1 is in an open state. State-7 has a post-powerstroke linker that is straight and stably docked in what we previously described as “post-2”(54) (in which the linker is upraised toward AAA3-AAA4), and a “kinked” buttress that has been attributed to MT-binding(23). The AAA1 pocket and AAA+ ring conformation for this state are similar to dynein structures with high MT-binding affinities in previous (PDB: 7z8g(8), 8fcy(23), 4w8f(44), 8pqv(22)) and this work. These observations suggest that state-7 represents a conformation primed for MT-binding, or that the high MT-binding affinity state can be intrinsically adopted by the motor even in the absence of MTs. States-5 and -6 are two novel states that have intermediate open AAA1 pockets and overall straight but undocked linkers, which likely represent intermediates during the powerstoke. The buttress in these two states exhibits an intermediate kink coupled with a flexible stalk-CC density, likely reflecting an intermediate MT-binding affinity. The stalk-CC is likely undergoing helix sliding(41) or local melting(76) in these two states, adopting a semi-α(62) or γ registry(53). Comparison of these three states (−5, -6, and -7) to a previously reported ADP dynein-1 structure (PDB:3vkg(47)) reveals close alignment of the AAA1 pocket and linker position with state-7. However, the buttress-stalk conformation of the 3vkg dynein is most similar to the state-6, suggesting that 3vkg represents a further intermediate state between states-6 and -7.

#### State-8

State-8, which has a nucleotide-free AAA1 pocket, is transient in excess ATP, but can be obtained in the apo condition. The large and small subunits of AAA1 are maximally separated in a wide-open state that permits ADP release (**Fig. 2d**), similar to a previous yeast apo motor structure (PDB:4aki)(48) and OAD-MT structures (PDB: 7k58,7k5b(54)). AAA3 has ADP bound even in apo condition. The linker is docked in the canonical post-powerstroke “post-1” position, and the buttress-stalk is in a high MT-binding affinity conformation.

Taken together, the range of conformational states we have observed for the full-length dynein-1 “in reaction” reflects not only the conformations observed for dynein structures solved in various nucleotide conditions but also novel intermediates never seen before (**Table 1**). Importantly, we observed that the linker, in addition to the previously described bent/docked conformation (pre-powerstroke) or straight/docked conformation (post-powerstroke), can adopt a straight but undocked conformation, likely representing crucial intermediate states during the powerstroke. These novel states demonstrate the importance of our approach and reveal novel insight into the mechanochemistry of the full-length dynein complex.

### The effect of nucleotide conditions on the conformational distributions of full-length dynein

Given the range of conformations we observed for the dynein motor, we wondered whether incubating dynein with different nucleotide analogs would lock the motor in a single conformation, as has been noted in prior cryo-EM and crystallographic studies(8, 23, 44, 47–49, 54, 77). To test this, we incubated full-length dynein (purified in 0.1 mM ATP) with or without additional analogs (i.e., 5 mM AMPPNP, ADP, ATP-Vi, or apo) and performed cryo-EM classification (**Extended Data Figs. 4, 5 and Supplementary Table 3–7**). To our surprise, we found that multiple conformational states are present in each condition (**Fig. 2e and Extended Data Fig. 6**). This may be due to the ability of dynein to adopt phi in all conditions(15); upon exiting phi with an ADP-bound to AAA1, which likely occurs stochastically, the dynein motors can then progress to various states before ADP release, and subsequent binding to the various nucleotide analogs. Below we describe the details of the conformations in each condition, as well as the observed nucleotide state (**Fig. 2e, f**).

#### ATP-Vi

It has been reported that dynein-2 adopts a closed AAA1 and a bent linker in the presence of ATP-vanadate (ATP-Vi, a mimic for ADP-Pi)(49). In our study of full-length dynein-1, we consistently observe closed AAA1 pockets. However, the linker displays both bent (states-1, 2) and straight (state-4) conformations in spite of the closed AAA1 pocket, aligning with previous observations of dynein-c(52, 78). The closed AAA1 pockets all exhibit density for ADP-Vi-Mg^2+^, in agreement with prior studies(23, 49). This suggests that the phi-motor (state-1), despite being in an autoinhibited state, retains the ability to hydrolyze ATP and release Pi, with the subsequent vanadate substitution occurring afterward.

#### AMPPNP

The non-hydrolyzable ATP analog AMPPNP has been widely used to trap cytoskeletal motors in an ATP-bound state(79, 80). In our experiments with AMPPNP, we observe the coexistence of multiple states. Surprisingly, AMPPNP does not bind to the AAA1 pocket in any of these states. Instead, the closed AAA1 pockets in both states-1 and -2 show clear density for ADP rather than ADP-Mg^2+^. We hypothesize that the absence of Mg^2+^ in the pocket results from an excess of AMPPNP relative to Mg^2+^ in the buffer (5 mM AMPPNP with 2 mM Mg^2+^). Consequently, if AMPPNP does not occupy the AAA1 site, Mg^2+^ may be chelated by excess AMPPNP. To test this, we determine the structures using a buffer containing 5 mM AMPPNP and 5 mM Mg^2+^. Consistent with our speculation, we observe ADP-Mg^2+^ (but not AMPPNP-Mg^2+^) in the closed AAA1 pockets. These results suggest that ADP is bound to the closed AAA1 pocket with such a strong affinity that even excess AMPPNP is unable to displace it without the ring opening, while Pi and Mg^2+^ can freely enter and exit the closed pocket.

In contrast to AAA1, we find that AAA3 is bound to AMPPNP in state-7 (**Extended Data Fig. 4**). However, AAA3 remains bound to ADP in the other states (1, 2, 5, and 6). This observation suggests that AMPPNP specifically targets the AAA3 site at a particular stage (state-7) of the cycle and can prevent the subsequent binding of AMPPNP to AAA1 site.

#### ADP and apo

As expected, in ADP and apo conditions, the proportion of molecules with open AAA1 pockets increased, but we also observed states with closed AAA1 pockets. Of note, in the apo condition, the wide-open apo AAA1 pocket conformation (state-8), which was very transient in the presence of ATP, was observed and co-existed with closed AAA1 states. The increased proportion of states-7 and -8 in ADP and apo conditions also supports our sequence determination shown in Figure 2a. In both conditions, the closed AAA1 pockets (states-1 and 2) have the same ADP-Mg^2+^ density as dynein in 5 mM ATP, indicating that the AAA1 pocket of full-length dynein can adopt a closed conformation even with low concentrations of ATP.

Taken together, our findings demonstrate that incubation with a certain nucleotide analog does not necessarily lock full-length dynein in one conformation but rather changes its conformational distribution. The structures found in these nucleotide conditions, combined with those in the ATP condition, provide a more comprehensive understanding of the dynamic conformational changes during the dynein mechanochemical cycle.

### Identification of a “molecular backdoor” for phosphate release from AAA1

States-1, -2, -3, and -4 in 5 mM ATP all have closed AAA1 pockets and AAA+ ring conformations. However, the linker in each of these states ranges from pre-powerstroke to a variety of intermediate states during the powerstroke (**Fig. 2a**). We observed a similar range of linker states for those dyneins with a closed AAA1 pocket in the ATP-Vi and AMPPNP conditions (**Fig. 2e**). Given the proposed role of AAA1 in governing linker position(37, 42, 49, 52, 78), these observations suggest there may be structural differences in AAA1 that are not apparent from a simple large-scale assessment of the open or closed state of this pocket. We thus more closely investigated whether there exist any potential small-scale changes in AAA1 among the different states. This revealed that the sensor-I loop adopts multiple distinct conformations in the presence of ATP (**Fig. 3a**). This loop is essential for the ATPase activity of AAA+ proteins(81) and has been proposed to facilitate Pi release via a backdoor mechanism(35). Notably, this loop adopts the same conformation in dyneins with closed AAA1 pockets (states-1, -2, -4) incubated in ATP-Vi (or 5 mM AMPPNP with 2 mM Mg^2+^) (**Fig. 3a**). However, the sensor-I loop becomes completely disordered when the AAA1 pocket is open (**Fig. 3b**), such as in states-7 and -8.

**Fig. 3.**
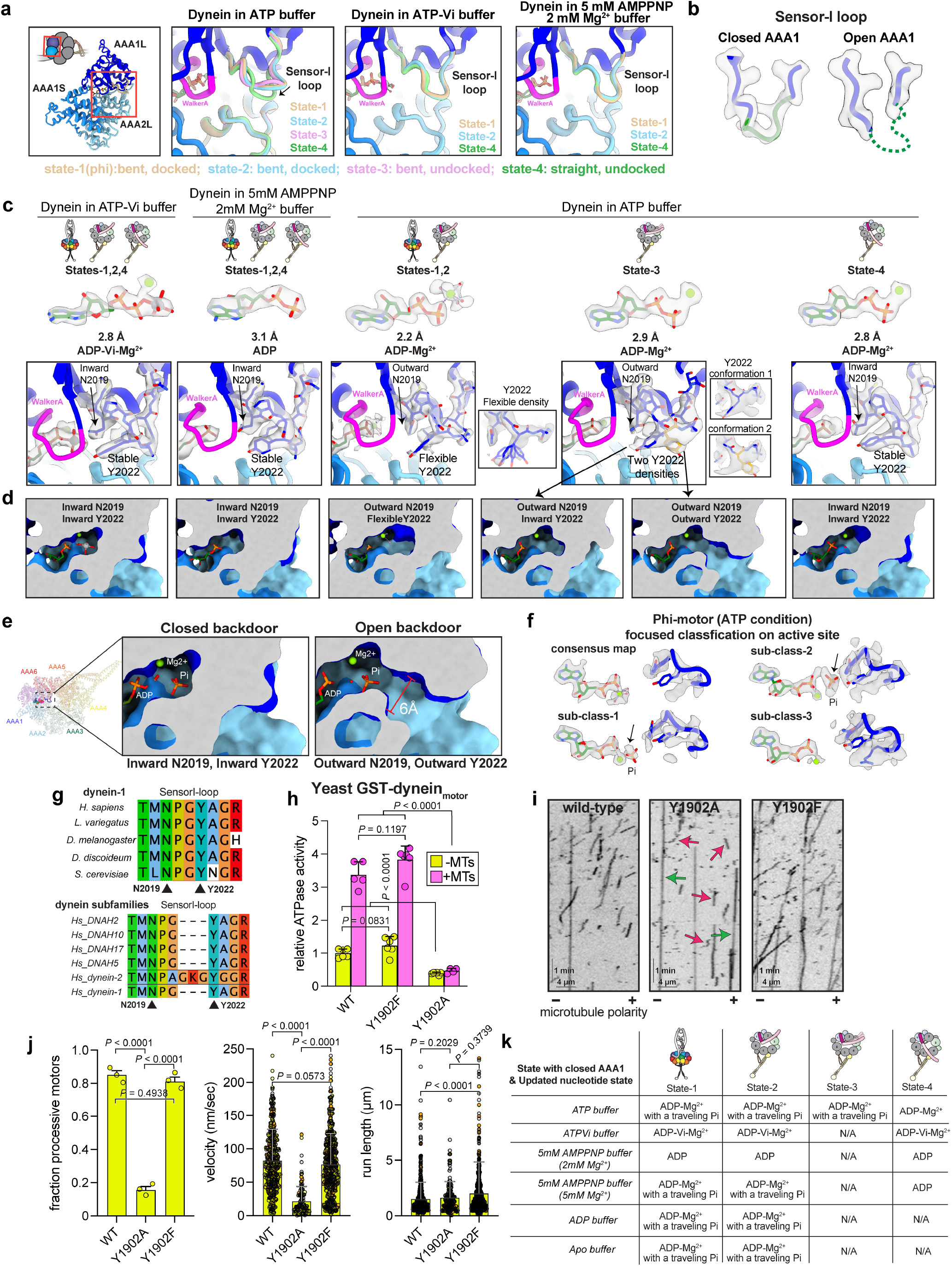
The release of Pi from AAA1 through a molecular backdoor mechanism. **(a)** Comparison of molecular models of closed AAA1 pockets in different states and nucleotide conditions. The sensor-I loop displays conformational heterogeneity when dynein is in ATP condition but not in ATP-Vi and AMPPNP conditions. **(b)** Cryo-EM densities of sensor-I loop in closed and open AAA1 pockets, showing that this loop becomes disordered when the AAA1 pocket is in open states. **(c)**) Structural comparison of the bound nucleotide and the sensor-I loop in closed AAA1 states. Transparent cryo-EM densities of the sensor-I loop are displayed in isosurface mode. The conformations of two residues, N2019 and Y2022, are highlighted. **(d)** A slice through the structures of the AAA1 pocket near the nucleotide-binding site, shown in surface representation.**(e)** Backdoor model for Pi release from dynein AAA1 controlled by N2019 and Y2022. **(f)** High-resolution 3D classification of the active site in the phi motor, showing the potential Pi group in various states of escape. **(g)** Conservation analysis of the sensor-I loop for dynein-1 in different species and dynein in different subfamilies (DNAH, dynein axonemal heavy chain; dynein-2, intraflagellar transport dynein). **(h)** Plot (mean ± SD, along with individual data points) depicting relative ATPase rates (in the absence or presence of 2 µM microtubules) for artificially dimerized yeast dynein motor domain fragments (from left to right, n = 6/5, 6/4, and 6/6 independent replicates from assays minus/plus MTs). Y1902 in yeast dynein corresponds to Y2022 in human dynein. **(i and j)** Representative kymographs (i) and plots (i; mean ± SD, along with individual data points) depicting indicated motility parameters determined from single-molecule motility assay (red arrows, diffusive motors; green arrows, processive motors; from left to right, n = 508/597, n = 148/880, and n = 679/837 processive/non-processive dynein molecules from three independent experiments). Microtubule polarity was determined with a wild-type dynein fragment (see Methods). P values in panel h were calculated using an ordinary one-way ANOVA followed by Tukey’s multiple comparison test. P values for panel j were calculated using a Kruskal-Wallis ANOVA followed by Dunn’s multiple comparison (for run length), or an ANOVA with Dunnett’s multiple comparison test (for velocity). **(k)** Summary of the updated nucleotide state among different closed AAA1 conformations. Note that the presence of a traveling Pi is based on the sensor-I loop conformations.

Close inspection of the cryo-EM densities of the sensor-I loop reveals two residues within this loop, N2019 and Y2022, that exhibit significant conformational changes (**Fig. 3c**). In both ATP-Vi and AMPPNP (low Mg^2+^) conditions, these two residues consistently orient towards the AAA1 active site and make direct contact with the AAA1 Walker-A motif and AAA2L. In the presence of ATP, these residues exhibit distinct conformations: in both states-1 and -2, N2019 points away from the active site (“outward”), while Y2022 adopts multiple conformations (“flexible density”). In state-3, which is absent in ATP-Vi and AMPPNP conditions, N2019 also points away from the pocket, and Y2022 has two alternative conformations. In state-4, the entire sensor-I loop exhibits stable density. Thus, despite the closed AAA1 pocket for all these states, there indeed exist small-scale, residue-level changes that very likely coordinate dynein mechanochemistry.

To investigate the role of the dynamics of these sensor-I loop residues during the ATPase cycle, we compared the local environment among the different states (**Fig. 3d**). This revealed that when both N2019 and Y2022 point outward, a channel appears to form, creating a possible escape route for Pi release. This observation supports the notion of a “molecular backdoor”(35) in which the coordinated rearrangement of residues in the sensor-I loop can establish a closed or open backdoor during the Pi release process (**Fig. 3e and Supplementary Video 3**). To further test this model, we performed focused classification of the AAA1 active site for the phi-motor (state-1) in ATP and captured what may be Pi at two different points during its release (**Fig. 3f;** see arrows). Importantly, the residues at the sensor-I loop, including N2019 and Y2022, are highly conserved between dynein-1 in different species and among dyneins (e.g., axonemal dynein and dynein-2; **Fig. 3g**), suggesting they are likely important for some aspect of dynein mechanochemistry.

The dynamic nature of Y2022 suggests it plays a crucial role in mediating the release of Pi. To test the importance of this residue in dynein function, we utilized a well-characterized GST-dimerized yeast dynein motor construct(43) and mutated the corresponding residue (Y1902) to either phenylalanine or alanine. Compared to the wild-type, Y1902F exhibits similar basal and MT-stimulated ATPase levels while Y1902A was defective in both (**Fig. 3h**). The addition of MTs only marginally stimulated the ATPase activity of the Y1902A mutant (by 20%, compared to 3.4- and 3.1-fold for wild-type and Y1902F, respectively), suggesting MT-binding is uncoupled from ATPase activity. Unlike human dynein, this minimal GST-dynein fragment is processive without the addition of dynactin or adaptor proteins, permitting assessment of motility without complicating factors. Single-molecule assays revealed that, unlike Y1902F, the Y1902A mutant exhibits significant motility defects, as apparent from the reduced fraction of active motors and the much lower velocity for those motors that exhibit processive runs (**Fig. 3i, j**). Intriguingly, a large proportion of the Y1902A motors (84.4%) appear to exhibit bidirectional diffusive behavior along MTs (see red arrows in kymographs), which is further suggestive of uncoupled MT-binding and AT-Pase activity. These functional data demonstrate the importance of this tyrosine in the catalytic and motility behaviors of dynein.

In light of this new backdoor model for Pi release, we can now more accurately assign the nucleotide state in different closed AAA1 pockets. Although we do not observe stable Pi density within the pocket, we can infer whether the closed pockets contain a traveling Pi by examining the conformations of residues N2019 and Y2022 (**Fig. 3k and Extended Data Fig. 7**). This reassessment reveals that the AAA1 pocket for states-1 and -2 in 5 mM ATP likely possess ADP-Mg^2+^ with a traveling Pi.

### Pi release triggers linker straightening during the active cycle

The backdoor Pi release mechanism prompted us to revisit the ATP turnover of the phi and open dynein. Our data indicate that phi dynein is indeed capable of hydrolyzing ATP, as apparent from the clear density of ADP-Mg^2+^, as well as by the presence of Vi in the ATP-Vi conditions (**Fig. 2b**). Furthermore, the dynamic sensor-I loop conformation suggests that a Pi is in the process of escaping the pocket (**Fig. 3**). However, due to the autoinhibited state of the motor domains, ATP hydrolysis and subsequent Pi release do not lead to a change in linker conformation (**Fig. 4a**). Similar to phi dynein, states-2 and -3 of open, uninhibited dynein also have dynamic sensor-I loop conformations that are indicative of a traveling Pi. Unlike phi dynein, the linker in state-3 shows a transitional conformation between bent and straight, suggesting a correlation between the sensor-I loop and linker position during Pi release. After the Pi is released in state-4 via the backdoor mechanism, as indicated by the stable densities for N2019 and Y2022, the linker adopts a straight conformation. This is consistent with previous kinetic measurements that the linker adopts a straight conformation after the release of Pi(42). Based on the Pi-free but closed state of the AAA1 pocket in state-4, we posit that subsequent to Pi release, and not prior to, the AAA1 pocket can adopt an open state.

**Fig. 4.**
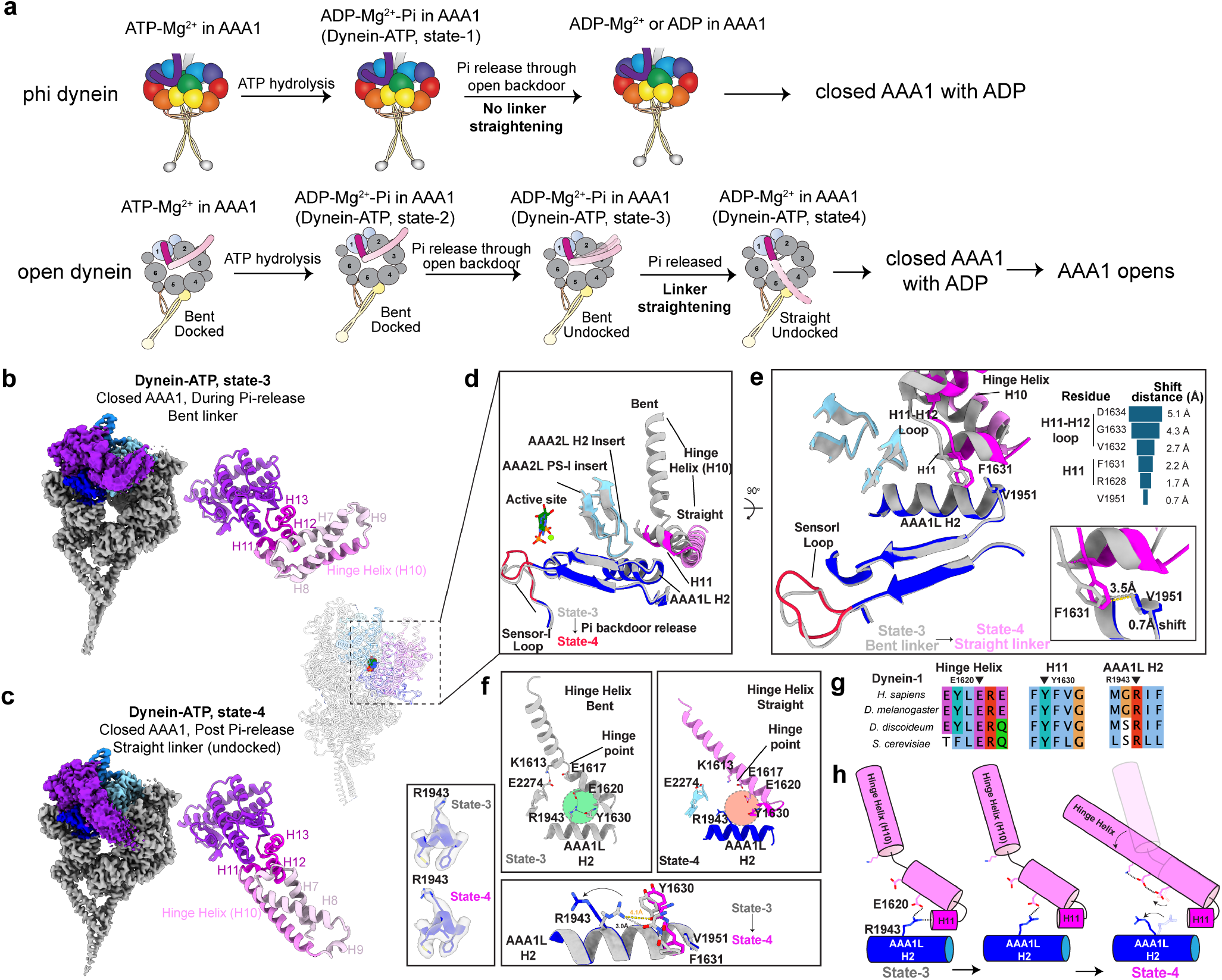
A mechanism for linker straightening triggered by the Pi release during the active cycle. **(a)** Cartoon models showing the process for Pi release for phi and open dynein. Unlike autoinhibited phi dynein, the release of Pi in open dynein results in the linker straightening. **(b**,**c)** Cryo-EM maps of states-4 and -3 with accompanying molecular models of the linker. The hinge helix in the linker is indicated. **(d)** Structural comparison of the state-3 (grey) and state-4 (color by domains). The AAA+ ring between the two states shares the same conformation while the sensor-I loop and linker adopt different conformations.**(e)** Linker-ring interface remodeling from state-3 (grey) to state-4 (colors). From state-3 to -4, H11, and the H11-H12 loop shift away from the AAA1L-H2, AAA2L PS-I insert and AAA2L H2 insert. Distance change values for the indicated residues are included. **(f-h)** Mechanism of the hinge helix straightening from state-3 to -4. In state-3, the bent hinge helix is stabilized by the conserved salt bridge and hydrogen bond interactions between R1943 in AAA2L-H2 and the hinge helix (f, g). During the linker-ring remodeling, R1943 flips away from the hinge helix in state-4, breaking these interactions which leads to the straightening of the hinge helix (f). The cartoon models depicting this process are shown on the right (h).

Whether the linker is in a bent or straight state is determined by the hinge helix (**Fig. 4b,c**)(49). However, the precise mechanism that triggers the straightening of the hinge helix is unclear. Our structures reveal that the release of Pi from AAA1 is coordinated with angstrom-scale, but significant conformational changes of the sensor-I loop (**Fig. 3**). These changes could theoretically be propagated to the linker-AAA+ ring interface via AAA1L’s central β-sheets, leading to large nanometer-scale conformational changes of the hinge helix, and subsequent linker straightening at an even larger scale (**Fig. 4d**). Close inspection of the linker-ring interfaces in states-3 and -4 reveal that the short linker helix 11 (H11) and the H11-H12 loop exhibit 1.7-5.1 Å displacements from state-3 to -4 (**Fig. 4e)**. These movements remodel the interactions between the linker and AAA1L H2, AAA2L PS-I insert, and AAA2L H2 insert. One notable change is the apparent disruption of a salt bridge (R1943-E1620) between states-3 and -4 (**Fig. 4f**). This salt-bridge pair is conserved across dynein species and was also reported in the dynein-2 structure(49) (**Fig. 4g**), suggesting it likely plays a crucial role in maintaining a bent linker. In state-3, this salt bridge is further stabilized by a hydrogen bond between R1943 and Y1630 in linker H11. In state-4, as linker H11 shifts away from the AAA1L H2, the hydrogen bond and salt bridge are both broken, causing the R1943 side chain to flip. Additionally, the AAA2L PS-I insert and H2-insert no longer contact the linker in state-4. The loss of all these interactions relieves the bent hinge helix from its distorted and restrained state(49, 52), thereby triggering the linker straightening (**Fig. 4f, h**).

### The communication between AAA1 pocket and the MTBD

It has been shown that AAA1 pocket dynamics regulate the affinity of the MTBD for MTs via registry changes in the stalk-CC(35). However, the precise mechanism for this long-range (24 nm) communication is unclear. Our extensive collection of dynein’s structural states reveals detailed pathways for the signal propagation from AAA1 to the MTBD (**Supplementary Video 4**). From state-4 to -5, the AAA1 pocket opens slightly, resulting in a movement of AAA2L towards AAA2S, and a small movement of AAA1L, which triggers the outward rotational movement of AAA6 (**Fig. 5a, b**). This further induces the CTD to swing from AAA3 to AAA5, and a consequent rotation of AAA5 causes the buttress to twist and the stalk-CC to adopt a “semi-α” registry, which is indicative of the MTBD adopting a higher MT-binding affinity. When the AAA1 pocket further opens from state-5 to -6 (**Fig. 2d**), AAA2L completely loses its interactions with AAA1L and moves towards AAA3 (**Fig. 5b**). This propagated signal appears to stop at AAA3L (**Fig. 5b**, note lack of movement beyond AAA3L in vector diagram), thus explaining the lack of change in the buttress-stalk conformation between state-5 and -6.

**Fig. 5.**
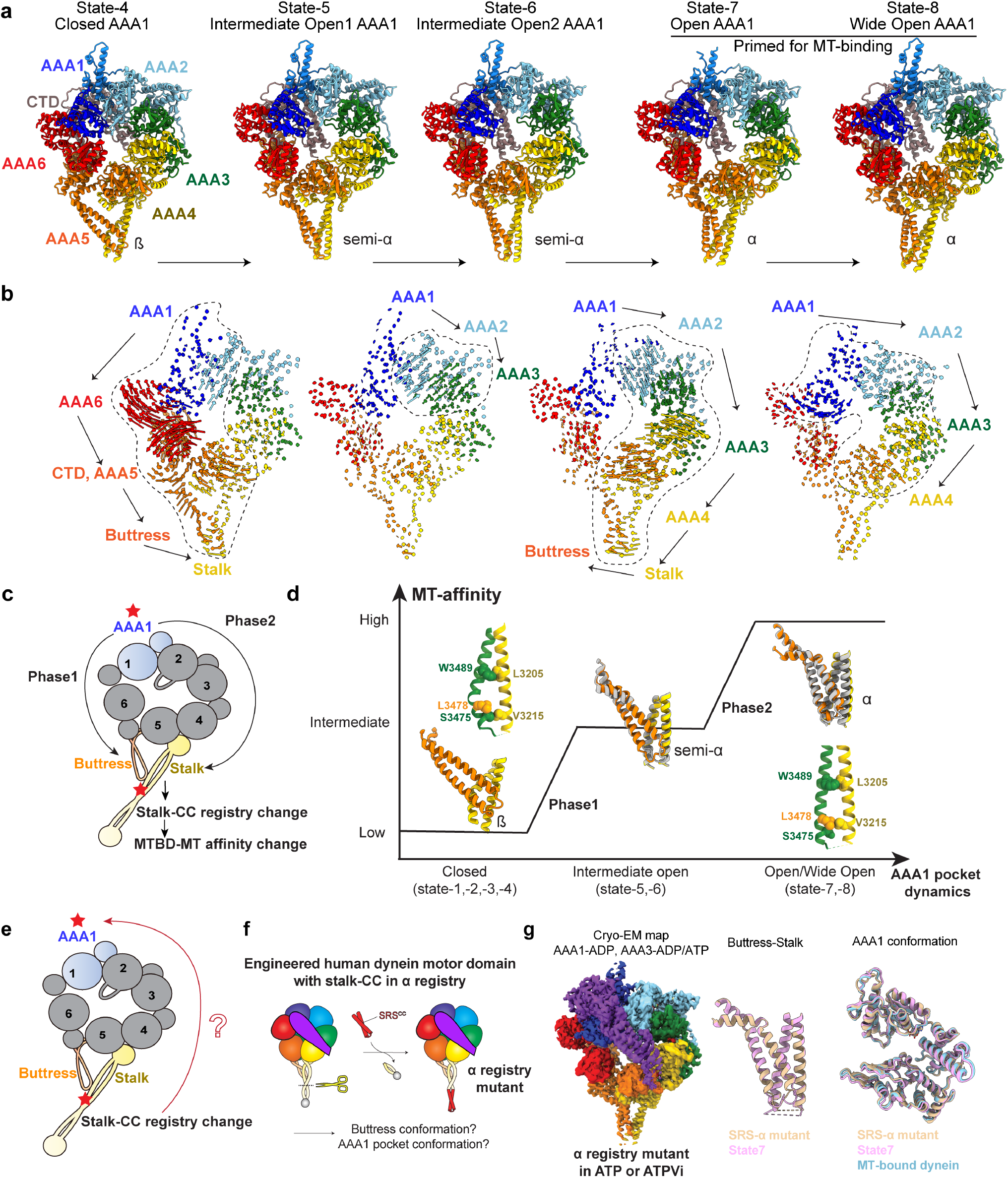
The mechanism for two-way communication between AAA1 and the MTBD. **(a)** Cartoon models illustrate the overall motor ring conformations from different states, each with varying degrees of the AAA1 pocket opening. The linker domain is hidden for better visibility of the motor ring states. **(b)** Adjacent models were aligned using the entire motor domain, and then vector maps were generated that depict pairwise alpha carbon interatomic distances (for every 3rd residue) between the AAA domains of adjacent states. The length of the lines is proportional to the calculated interatomic distances. The major movements and directions are highlighted by dashed areas and arrows. **(c)** The cartoon model illustrates the signal propagation directions from AAA1 to the stalk-CC in two distinct phases. **(d)** Changes in dynein-MT binding affinity (Y-axis), as assessed by buttress-stalk conformations, as a consequence of AAA1 pocket conformations in the different states (X-axis). **(e)** Cartoon model illustrating the communication from the MTBD to AAA1 via the stalk-CC. **(f)** Cartoon model depicting the design of the α registry-locked dynein motor domain mutant. **(g)**) Cryo-EM reconstructions of the α registry mutant in ATP or ATP-Vi conditions. In both conditions, the motor domains exhibit only open AAA1 pockets. As expected, the buttress-stalk conformation is consistent with a high MT-binding affinity.

Upon transition to state-7, the AAA2L domain continues to move away from AAA1L, but in a different direction: toward AAA3L **(Fig. 5b**). This translation is propagated to the buttress-stalk via AAA4L and AAA5L, leading to the adoption of the α-registry by the stalk-CC. From state-7 to -8, the AAA1 pocket opens completely, with a sufficient separation between AAA1L and AAA1S that permits ADP release. This pocket widening slightly affects the conformations of AAA2-5L but does not affect the buttress-stalk conformation (**Fig. 5b**). Our results strongly support a model in which AAA1 conformational signals are propagated in two phases (**Fig. 5c, d**): first through AAA6, CTD, AAA5, and the buttress to stalk-CC (to adopt the semi-α registry), and then subsequently via AAA2, AAA3, and AAA4 to complete the adoption of the α registry by the stalk-CC.

Our results show how AAA1 transmits its conformational signals to the buttress-stalk region to change the stalk-CC registry to affect MTBD-MT affinity. However, previous studies have shown that locking the stalk-CC in different registries affects the ATPase activity, motility, and tension-sensing of dynein motors(41, 53), supporting a model in which the MTBD can also transmit signals back to the AAA+ ring through the stalk-CC (**Fig. 5e**). To investigate the structural basis of the reverse communication, we performed cryo-EM analysis of a human dynein motor domain mutant in which the stalk-CC is locked in the α registry(23) (to mimic the MT-bound state; Fig. 5f, Supplementary Table 8). This revealed that in the presence of ATP, this mutant consistently has an open ADP-bound AAA1 pocket, and a kinked buttress conformation(23) (**Fig. 5g**), resembling that of WT full-length dynein in state-7. Even in the presence of ATP-Vi, the AAA1 pocket is bound to ADP and remains in the same open conformation, consistent with this mutant not undergoing Vi-dependent photocleavage (a readout for Vi binding)(23). These results suggest that locking the stalk-CC in the α-registry forces the AAA1 pocket to adopt an open conformation. Taken together, we provide a structural basis for a comprehensive understanding of the two-way communication between AAA1 and the MTBD.

### Structures of MT-bound dynein motors reveal that MTs accelerate ADP release

To gain insight into the role of MTs during dynein’s mechanochemical cycle, we obtained four distinct structures for MT-bound dynein motor domains (**Fig. 6a**), with AAA1 in either apo or ADP-bound states, all of which possess stably docked linkers and kinked buttress. Exclusion of nucleotide from the buffer resulted in AAA1 being in an apo state, AAA3 and AAA4 being bound to ADP, and the linker being in the canonical “post-1” post-powerstroke state. In ADP-containing buffer conditions, we identified ADP in AAA1, AAA3, and AAA4, with the linker docking at one of two different positions (post-1 and post-2). In the presence of AMPPNP, both AAA3 and AAA4 are bound to AMPPNP, while AAA1 remains ADP-bound, and the linker is docked at only post-1.

**Fig. 6.**
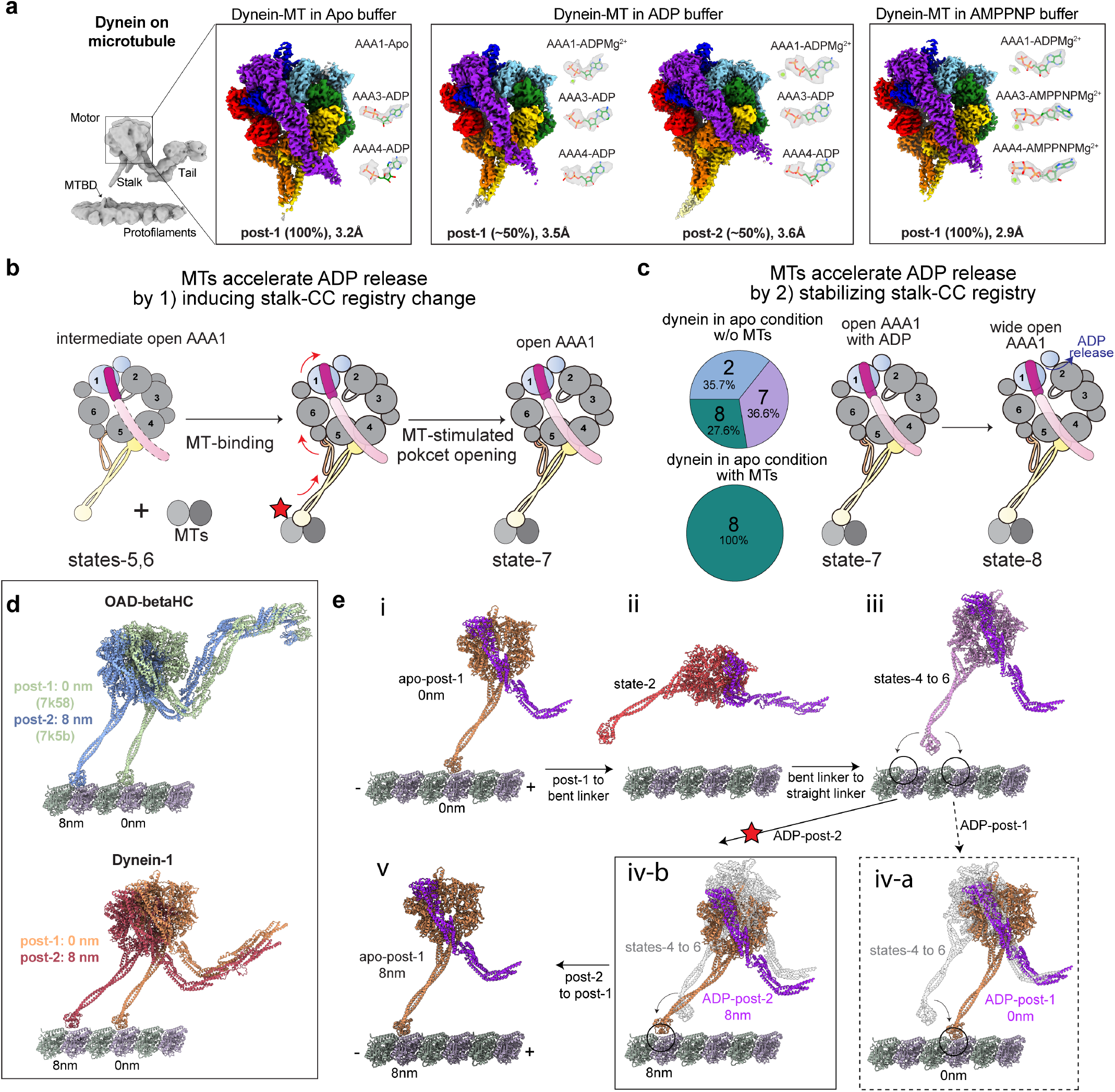
The mechanism for ATPase stimulation by MTs and dynein’s unidirectional stepping along MTs. **(a)** High-resolution cryo-EM maps of MT-bound dynein motor domains in different nucleotide conditions. **(b)** The pie chart of the distribution of various states of dynein in active cycle with ATP. The Pi release and ADP release steps correlated with different states are shown. **(c)** The cartoon model depicts the mechanism of dynein mechanochemical cycle acceleration by MTs. MT-binding rapidly induces the stalk-CC registry from semi-α to α registry, therefore forcing the AAA1 pocket to adopt an open state (left). MT-binding, by stabilizing the stalk-CC registry, allows the further wide opening of the AAA1 pocket and leads to the final ADP release (right). **(d)** Dynein adopting post-2 linker docking mode intrinsically favors a forward step, based on the previous OAD-MT structures (PDB: 7k58, 7k5b(54)). Aligning dyneins by their N-termini (to the right of the models in the picture) show that the post-2 dynein motor for both OAD and dynein-1 adopts a forward stepping pattern with respect to the post-1 dyneins.**(e)** Proposed mechanism for dynein unidirectional stepping along MTs. **(i)** Dynein is bound to MTs in the apo state in post-1 mode (after completing a step). **(ii)** Upon ATP binding and/or hydrolysis, dynein unbinds MTs with a bent linker. **(iii)** Upon Pi release from AAA1, dynein with a straight but undocked linker (states-4 to 6) with low and intermediate MT-binding affinity. **(iv-b)** The preferred post-2 mode in ATP conditions (denoted by a star) creates a net bias towards the minus end of MTs and allows MTBD to take an 8 nm forward step. Note that adopting a post-1 mode upon MT-rebinding **(iv-a)** leads to a 0 nm step. The cartoon model from state-4 is shown in grey. **(v)** Switching from a post-2 to a post-1 mode and/or subsequent ADP release leads to an 8 nm forward movement of the entire dynein complex.

To assess the impact of MT binding on dynein motors, we compared these structures with the MT-unbound structures described above (**Extended Data Fig. 8**). Strikingly, three of the MT-bound motor domain structures (i.e., those from apo and ADP conditions) matched very well with structures from the conformational landscape of MT-unbound dynein (Cα RMSD < 0.3 Å with state-7 or -8). In contrast, MT-bound dynein incubated in AMPPNP-containing buffer exhibits a 2.3 Å-5.1 Å Cα RMSD in comparison to the MT-unbound motors, with the closest match being to state-These findings indicate that the dynein motor domain can naturally sample the MT-bound state even in the absence of MTs. However, it is noteworthy that the MT-bound motors exhibit more stable cryo-EM densities in the buttress-stalk and linker regions compared to their MT-unbound counterparts, suggesting MT-binding can further coordinate the buttress-stalk conformational change and linker docking during the dynein stepping cycle(53). In addition, the states-5 and -6 structures with intermediate open pockets are no longer found in MT-bound motors, providing additional evidence that MT binding indeed stabilizes the stalk-CC registry, the AAA+ ring conformation, and the linker-AAA docking, all of which is likely triggered via the MTBD-MT contact.

How might MTs accelerate the ATPase activity based on these results? It has been proposed that ADP release is the rate-limiting step for dynein(82) and that MTs accelerate this step(70). In agreement with this model, we find that approximately 45% of MT-unbound dynein motors exhibit ADP-bound intermediate AAA1 open pockets with their stalk-CC in the semi-α registry (states-5 and -6; Fig. 2a). MT-binding shifts these two intermediates to state-7 by inducing the stalk-CC to adopt the α registry(62) (**Fig. 6b**), which leads to further opening of the motor ring and the AAA1 pocket for subsequent ADP release. MT-binding can also induce the adoption of the “wide open” state of the AAA1 pocket, as observed in state-8, which allows ADP release. In support of this, nucleotide-free buffer conditions lead to the identification of both states-7 (i.e., ADP-bound AAA1) and -8 (i.e., apo AAA1) for MT-unbound dynein, but only lead to the presence of state-8 for MT-bound dynein (**Fig. 6c**). These results explain that two limiting steps, stalk-CC registry change(69), and ADP release(82), are strictly coupled and concurrently stimulated by MT-binding. In addition, our evidence for a Pi backdoor release mechanism in low-MT binding affinity states supports a previous model that MT-binding does not affect the Pi release step(70).

### Post-2 linker-ring docking mode provides a net bias for dynein minus-end stepping

The linker swing vector model has been proposed to explain the mechanism of dynein unidirectional motility(51, 58, 83). However, which step during the mechanochemical cycle provides a net minus end-directed bias remains unknown. We analyzed the different linker-ring interaction interfaces (**Extended Data Fig. 9a**) and found that the two different linker docking modes (post-1 and post-2) may directly impact dynein stepping behavior, similar to previous reports on OAD(54). The observation of a post-2 linker position in dynein-1 indicates that this docking mode is conserved (**Extended Data Fig. 9b-e**).

Based on previous OAD-MT structures, it was proposed that dynein in post-2 mode would position its MTBD 8 nm forward of dynein in post-1 mode (**Fig. 6d**). To investigate the role of the two different docking modes in dynein stepping, we assessed how the linker position would correlate with dynein stepping behavior (**Fig. 6e and Supplementary Video 5**). When AAA1 is in the apo state, the dynein motor binds to MTs in post-1, which is the starting position (0 nm; Fig. 6ei). ATP binding and/or hydrolysis at AAA1 leads to the adoption of the pre-powerstroke state, and the detachment of dynein from MTs (due to the low affinity for MTs in this state; Fig. 6eii). The Pi backdoor release and linker straightening process likely induce the AAA+ ring to rotate relative to the tail rather than pull the tail region forward because the dynein is in the low MT-binding affinity state(83) (**Fig. 6eiii**). After subsequent AAA1 pocket-opening, the motor can rebind to MTs with an ADP-bound open pocket. If the motor domain adopts the post-1 mode, the MTBD returns to its original position without any forward stepping (**Fig. 6eiv-a**). By contrast, stepping with the motor domain in post-2 mode generates a net bias towards the minus-end of MTs, allowing the dynein MTBD to bind 8 nm forward along the MT (**Fig. 6eiv-b**). In support of this idea, the MT-unbound motors in state-7 (ADP-bound open AAA1) primarily adopt the post-2 mode in the presence of ATP (**Fig. 2a**). Finally, the transition from ADP-post-2 to ADP-post-1 – or to apo-post-1 – results in further translocation (total 8 nm) of the cargo-binding tail domain (**Fig. 6ev**).

### Crosstalk between AAA1 and AAA3 revealed by structural survey of motor domains

In our large-scale collection of dynein structural states, we observe that the regulatory site AAA3 alternates between closed (ATP or AMPPNP-bound) and open (ADP-bound) conformations. To explore the communication between AAA1 and AAA3, we conducted a structural survey (**Extended Data Fig. 10a**) using all available dynein-1 motor domain structures from our work (43 PDBs) and previous studies (13 PDBs). For each PDB, we extracted the following information: the conformation and nucleotide state of the AAA1 and AAA3 pockets, the linker conformation, and the motor’s affinity for MTs. We then performed statistical analysis to assess the impact of AAA1 on AAA3, the linker, and MTBD affinity, and vice versa (Table 2, Extended Data Fig. 10b, c).

We found that in the closed AAA1 state (21/56 structures), AAA3 is always open. Among these 21 structures, AAA3 is always ADP-bound, with the exception of one structure (PDB: 7mgm(77)) in which ATP is bound to AAA3 due to a Walker-B mutation. Conversely, in the closed AAA3 state (14/56 structures), AAA1 is either in an intermediate open or fully open state. In all of these cases, AAA1 is ADP-bound, except for PDB 4w8f(44) (AMPPNP in AAA1 due to a Walker-B mutation). These two exceptions also show that the desired behavior of Walker-B mutation to induce pocket closure can be overrode by the other pocket’s conformation. When AAA1 is in intermediate open or open states, AAA3 exhibits dynamic behavior, and vice versa.

These observations suggest that the closed AAA1 state inhibits ADP release in AAA3, and similarly, the closed AAA3 state inhibits ADP release in AAA1 (**Extended Data Fig. 10d**). This mutual inhibition model explains why ATP or AMPPNP-bound AAA3 is only observed in specific states (states-5 to -7). Furthermore, when AAA3 is closed due to ATP or AMPPNP binding, it locks AAA1 in an open, ADP-bound state, resulting in a straight, docked linker and high MT-binding affinity, consistent with previous observations(8, 59, 84). We also find that, unlike open AAA1 pockets, which have a direct and specific impact on the conformation of the linker and the motor’s affinity for microtubules, open AAA3 pockets do not exert such direct control. This observation supports a previously proposed model in which AAA3 regulates the motor’s activity through a switching mechanism(59, 84).

## Discussion

### Technical advances to explore the dynein mechanochemical cycle

In this work, we demonstrate that our improved cryo-EM pipeline (**Fig. 1**) can identify the conformational landscape of full-length human dynein-1 in reaction with ATP. This approach contrasts with previous studies that employed truncations, mutations, and specific nucleotide conditions to lock dynein in one state(44, 47–49). Our pipeline not only captures nine distinct motor states but also resolves them at high resolutions (**Fig. 2**). The 2.2 Å structure of the phi motor domain provides a solid structural basis to analyze the molecular interactions at an atomic level, particularly the residues involved at the reactive center. Systematic analysis of the dynamic range of motor domain structures of full-length open dynein, both on and off MTs, permits an extremely comprehensive understanding of the mechanochemical cycle of dynein (**Fig. 2–6**). Our approach also resolves the MT-bound dynein motors at high resolutions, shedding light on the impacts of MT-binding on dynein (**Fig. 6**). The numerous new intermediate states reported in this work, which are technically challenging to acquire using conventional approaches, substantially expand our knowledge of the dynein conformational landscape.

### The release of Pi through a molecular backdoor mechanism

We provide structural and functional evidence that supports the existence of a “backdoor” for Pi release that involves small but significant rearrangements of the AAA1 sensor-I loop (**Fig. 3**). In the presence of ATP, two residues in the sensor-I loop – N2019 and Y2022 – are found in various conformations in different motor states. We speculate that N2019 senses the Pi cleavage while Y2022 gates the release of Pi. In support of this idea, N2019 closely interacts with the vanadate group when AAA1 is bound to ADP-Vi (2.6 Å). The cleavage of the ATP forces N2019 to adopt outward conformation to accommodate the Pi. Meanwhile, Y2022, which exhibits multiple conformations, creates a gated escape route for Pi release. It likely performs this function via its large aromatic side chain, since the Y1902F mutant was functional in ATPase and motility assays. When the Pi is fully released, both residues stably contact the Walker-A motif, completing the Pi release step and allowing the subsequent pocket opening processes.

Furthermore, our data strongly support a mechanism whereby the conformational changes in the sensor-I loop transmit Pi-release signals to the process of linker straightening (**Fig. 4**). However, this process is sterically restrained in the autoinhibited phi dynein in which Pi release does not lead to linker straightening. These findings also elucidate why the bent or straight linker are both found in the presence of ATP-Vi for human dynein-1 (this study), yeast dynein-1(44), dynein-2(49), and axonemal dynein-c (52, 78). As Pi is in the process of releasing through the backdoor, vanadate can attack the active site while the linker is in multiple intermediate states, therefore stabilizing dynein in a closed motor ring state with either a bent or straight linker.

### A new perspective of dynein conformation distribution under various conditions

Our findings also show that a specific nucleotide environment influences the conformational distribution of the motor domain for full-length dynein. A recent structure of the DDA-LIS1 complex bound to MTs in the presence of AMPPNP revealed a pre-powerstroke state motor domain bound to LIS1(22). In this motor-LIS1 complex with a bent linker, the AAA1 pocket binds ADP with a closed conformation, which is speculated to be induced by LIS1(22). Our data demonstrates that this motor state (state-2), without LIS1 binding, can be directly observed under ATP, ADP, AMPPNP, and even apo conditions in the absence of LIS1 (**Extended Data Fig. 4**), indicating that this is an intrinsic property of the dynein motor domain. However, we cannot rule out that the presence of LIS1 may stabilize and enrich for this state in the context of a MT-bound DDA complex(22). In the future, by comparing the conformational landscape of dynein with LIS1 or other regulators to the many structures reported in this study, the impacts of regulators on dynein can be better characterized.

### The powerstroke of dynein consists of multiple steps

The powerstroke of dynein involves the transition of the AAA+ ring from a closed (MT-unbound) to an open (MT-bound) conformation, coupled with the straightening of the linker from a bent to a straight conformation(33, 35, 85). This transition has been hypothesized to generate the force necessary for dynein to pull its cargo forward. The existence of state-4 in our dataset indicates that linker straightening occurs before pocket opening and even MT-binding (**Fig. 3**). We also identify two additional states (states-5, and -6) with partially open ADP-bound AAA1 pockets that appear to represent intermediates before the MT-bound state (**Fig. 4**). Note that all three states (4 through 6) feature straight (as apparent at low contour levels), but undocked linkers. Upon dynein binding to MTs, the straight linker is stably docked on the motor ring in one of the two modes (post-1 or post-2).

Based on the existence of these newly identified linker intermediates, we propose a new model for the force-generating powerstroke (**Supplementary Video 5**). We designate state-2, characterized by a closed AAA+ ring with an ADP-Pi bound AAA1 pocket (i.e., post-ATP hydrolysis and before Pi release), and a bent, stably docked linker, as the pre-powerstroke state, consistent with a previous structure(49). We propose that the powerstroke proceeds through the following steps: 1) Pi is released from AAA1 via the backdoor (while the AAA1 pocket and AAA+ ring remain closed), and the linker transitions from a bent, stably docked state to a straight, unstable conformation. Note the MT-binding affinity remains low, and dynein consequently unbinds from MTs. 2) The AAA+ ring transitions from closed to intermediate open states while the linker remains straight and undocked. 3) Upon MT-binding (and adoption of a high MT-binding affinity state), the linker transitions to a conformation in which it is docked in a post-2 state (the preferred state for dynein in the moments before MT-binding), coincident with the AAA+ ring transitioning to an open state. 4) The linker transitions to a post-1 state in the ADP-bound or apo AAA1 state, resulting in the translocation of cargo by 8 nm. In this model, the force necessary for dynein to pull its cargo is primarily generated during steps 3 and 4 as dynein is anchored on MTs.

### Model for full-length human dynein-1’s mechanochemical cycle

Our work significantly improves our understanding of dynein’s mechanochemical cycle (**Fig. 7**). We find that the motor domain from the autoinhibited phi dynein is competent for ATP hydrolysis and Pi release. However, the closed AAA1 pocket in phi dynein sterically hinders ADP release as the Pi release channel is not large enough to accommodate ADP release. The motors engaged in the mechanochemical cycle are predominantly in ADP-bound states, consistent with ADP-release being the limiting step, and with the fact that dynein only possesses basal ATPase levels off MTs(70, 82). By comparing the conformational landscapes for dynein on and off MTs, we propose the following model to explain its mechanochemical cycle with MTs. Note that dynein can still progress through this cycle when unbound from MTs (**Fig. 2**).

**Fig. 7.**
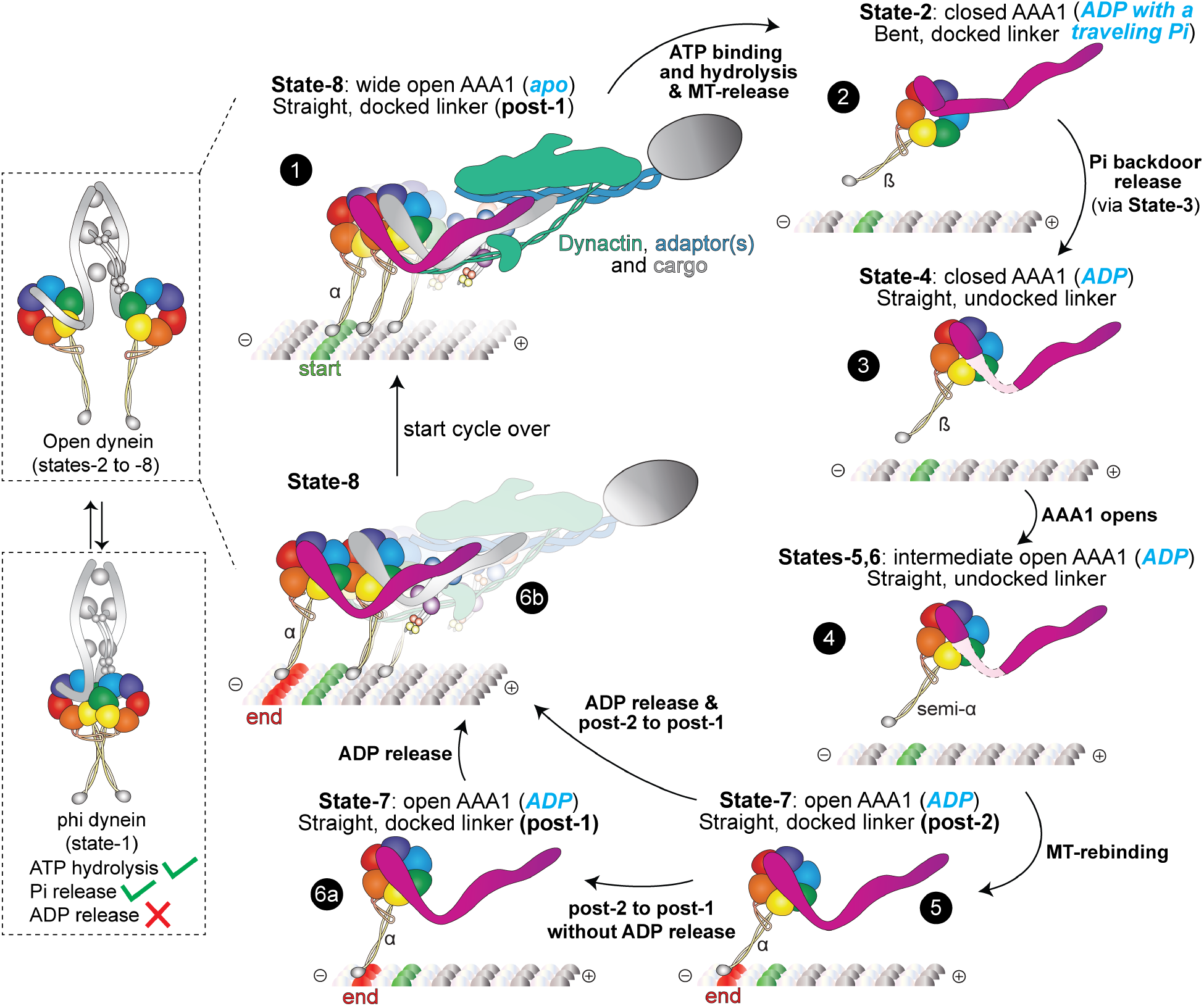
Model for the mechanochemical cycle of full-length human dynein-1. The full-length dynein adopts two autoinhibited states: state-0 in the open conformation, and state-1 in phi conformation. Motor domains in autoinhibited states and the active states exist in an equilibrium. The mechanochemical cycle of the active motor states starts with the apo AAA1 state. ATP binding and/or hydrolysis induces pocket closure and triggers the MT release. The linker changes from straight, stably docked to bent, stably docked conformation. The backdoor release of Pi triggers the straightening of the linker. The presence of MTs actively engages in the pocket-opening process by inducing the stalk-CC registry to change from semi-α and α states. The linker conformation changes from the straight, unstably docked to straight, stably docked conformation upon MT-binding. The linker-docking mode transitions and ADP release completes the cycle and dynein takes a forward step.

1) ATP binding to the apo AAA1 state (state-8) and/or subsequent hydrolysis induce pocket closure, triggering a registry change in the stalk-CC from α to β and a straight-to-bent shift in the linker(35). 2) Following ATP hydrolysis, the sensor-I loop facilitates Pi release from AAA1 via a molecular backdoor mechanism. After the release of Pi, the linker is straightened but not stably docked on the closed motor ring. 3) The ADP-bound AAA1 pocket gradually transitions from closed to intermediate-open and open states. Throughout this process, the buttress and stalk respond to the pocket-opening signal across the motor ring, resulting in semi-α and α registry stalk-CC configurations. MT-binding significantly accelerates this step by actively triggering the adoption of the α registry of the stalk-CC that eventually facilitates ADP release. 4) Further widening of the AAA1 pocket ensures ADP release and dynein thus pulls its cargo one step forward (see above), completing the current cycle. It is worth noting that our current model focuses exclusively on the primary AT-Pase site, AAA1. Although we have observed crosstalk between AAA1 and AAA3, our current cryo-EM analyses are not sufficient to definitively resolve the ongoing debate about whether the hydrolysis of two ATP molecules (e.g., at both AAA1 and AAA3) is required(86), or if only one ATP (at AAA1) is hydrolyzed during the complete cycle(87, 88).

Our model, supported by previous structural studies and over 40 high-resolution structures in this work, sheds new light on many long-standing questions in the field(35). Our approach to determining the conformational landscape of dynein “in reaction” – on and off MTs – has revealed valuable insight that was impossible with prior approaches that relied on strategies that aimed for conformational homogeneity. Our structures obtained with full-length dynein also provide opportunities to analyze potential coordination between the two motor domains off and on MTs that will be addressed in future studies. In addition to providing numerous testable hypotheses (via mutagenesis combined with functional assays), our high-resolution intermediate states of the dynein motor provide a strong structural basis for computational simulation(89) and protein engineering(90). Lastly, our work featuring high-resolution dynamic cryo-EM analysis provides a new paradigm that can be expanded to provide deeper mechanistic insight into the entire dynein transport machinery on MTs(2) (in complex with dynactin and a cargo adaptor), or regulation by its cofactors (e.g., LIS1(18), Ndel(24)).

## Acknowledgements

We are very grateful to members of the Zhang and Markus laboratories for their valuable discussions. We thank R. Yang for the initial dynein-1 expression and preparations in the lab. This work was funded by the NIH/NIGMS (R35GM139483 to S.M.M., and R35GM142959 to K.Z.) and in part by a Collaboration Development Award Program (to K.Z.) from the Pittsburgh Center for HIV Protein Interactions (U54AI170791). We would like to thank K. Zhou, J. Lin, M. Llaguno, and S. Wu, for their help with cryo-EM data collection at Yale Cryo-EM facility. The Yale Cryo-EM Resource is funded in part by the NIH grant S10OD023603 awarded to F. Sigworth. We thank L. Wang, J. Kaminsky, and G. Hu at the Laboratory for BioMolecular Structure (LBMS) for help with cryo-EM data collection. The LBMS is supported by the DOE Office of Biological and Environmental Research (KP1607011).

## Author contributions

K.Z. designed the study. J.Y., Y.W. purified full-length human proteins, and J.Y., Y.W., P.C. prepared the cryo-EM samples and collected the data. J.Y., Y.W., P.C., K.Z. processed the images, and built PDB models. S.M.M. purified the human MT-B dynein motor domain. I.C.G. generated yeast strains, and purified proteins from yeast. I.C.G. and S.M.M. performed and analyzed ATPase and single molecule assays. P.C., J.Y. Y.W., K.Z., and S.M.M. generated figures and movies. P.C., J.Y., and K.Z. wrote the manuscript. P.C., J.Y., Y.W., K.Z., and S.M.M. edited and revised the manuscript. S.M.M. and K.Z. acquired funding.

## Data availability

Models and cryo-EM maps have been deposited in the PDB and EMD as follows:

Dynein in ATP buffer: PDB-9BLY/EMD-44681 (phi particle dynein), PDB-9BLZ/EMD-44682 (state-1), PDB-9BM0/EMD-44683 (state-2), PDB-9BM1/EMD-44684 (state-3), PDB-9BM2/EMD-44685 (state-4), PDB-9BM3/EMD-44686 (state-5a), PDB-9BM4/EMD-44687 (state-5b), PDB-9BM5/EMD-44688 (state-6), PDB-9BM6/EMD-44689 (state-7a-post-2), PDB-9BM7/EMD-44690 (state-7b-post-2), PDB-9BM8/EMD-44691 (state-7c-post-2).

Dynein in ATP-Vi buffer: PDB-9BMF/EMD-44697 (state-1), PDB-9BMG/EMD-44698 (state-2), PDB-9BMH/EMD-44699 (state-4).

Dynein in 5 mM AMPPNP buffer with 2 mM Mg^2+^ : PDB-9BMJ/EMD-44701 (state-1), PDB-9BML/EMD-44702 (state-2), PDB-9BMM/EMD-44703 (state-4), PDB-9BMN/EMD-44704 (state-5), PDB-9BMO/EMD-44705 (state-6), PDB-9BMP/EMD-44706 (state-7-post-2).

Dynein in 5 mM AMPPNP buffer with 5 mM Mg^2+^ : PDB-9DH5/EMD-46856 (state-1), PDB-9DH6/EMD-46857 (state-2), PDB-9DH7/EMD-46858 (state-4), PDB-9DH8/EMD-46859 (state-5), PDB-9DH9/EMD-46860 (state-7-post-2), PDB-9DHA/EMD-46861 (state-7-post-1)

Dynein in ADP buffer: PDB-9BMR/EMD-44708 (state-1), PDB-9BMS/EMD-44709 (state-2), PDB-9BMT/EMD-44710 (state-5), PDB-9BMU/EMD-44711 (state-6), PDB-9BMV/EMD-44712 (state-7-post-1), PDB-9BMW/EMD-44714 (state-7-post-2).

Dynein in apo buffer: PDB-9BMY/EMD-44715 (state-1), PDB-9BMZ/EMD-44716 (state-2), PDB-9BN0/EMD-44717 (state-7-post-2), PDB-9BN1/EMD-44718 (state-8-post-1).

Dynein bound to microtubules: PDB-9BMA/EMD-44693 (apo-post-1), PDB-9BMB/EMD-44694 (ADP-post-1), PDB-9BMC/EMD-44695 (ADP-post-2), PDB-9BMD/EMD-44696 (AMPPNP-post-1).

Dynein motor domain alpha registry mutant in ATP buffer: PDB-9BN3/EMD-44720 (class-1), PDB-9BN4/EMD-44721 (class-2).

Dynein motor domain alpha registry mutant in ATP-Vi buffer: PDB-9BN5/EMD-44722 (class-1), PDB-9BN6/EMD-44723 (class-2).

## Methods

### Full-length human dynein-1 purification

The plasmid encoding full-length human dynein-1 was graciously provided by the Andrew Carter Lab (Addgene plasmid 11903). This plasmid featured the dynein-1 heavy chain fused to an N-terminus for the His-ZZ-SNAPf tag, followed by the Tobacco Etch Virus (TEV) protease cleavage site. Recombinant baculoviruses were produced in sf9 insect cells via transfection of isolated Bacmid obtained from DH10MultiBac competent cells (Geneva Biotech) using Cellfectin® II (Gibco). Expression of dynein-1 in sf9 cells was accomplished by infecting them with P2 virus at a cell density of 2.5 million cells/ml. 28 ml of P2 virus was added into a 1.4 L culture of sf9 cells. Following 75 hours of incubation, the infected cells were harvested by centrifugation at 1000 rcf for 15 minutes at 4 °C. The resulting cell pellet was flash-frozen in liquid nitrogen and stored at −80 °C until further use. For dynein purification, frozen cell pellets from 1.4 L were thawed and resuspended in 100 ml of lysis buffer (50 mM HEPES pH 7.2, 100 mM NaCl, 1 mM DTT, 0.1 mM ATP, 10% glycerol) supplemented with 2 protease inhibitor tablets (Roche) and 2 mM PMSF. The cell suspension was lysed using a 40 mL Dounce tissue grinder (Wheaton) with 15-25 strokes on ice, followed by clarification via ultracentrifugation at 65k rpm with a Ti70 rotor for 1 hour at 4 °C. The resulting supernatant was incubated with 3 mL of pre-equilibrated IgG Sepharose affinity beads (Cytiva) on a roller at 4 °C for 3 hours. Subsequently, the beads were packed into a gravity column, washed sequentially with 200 mL of lysis buffer, followed by 200 mL of TEV buffer (50 mM Tris–HCl pH 7.4, 148 mM K-acetate, 2 mM Mg-acetate, 1 mM EGTA, 10% glycerol, 0.1 mM ATP, 1 mM DTT). The beads were then transferred into a 15 ml tube filled with TEV buffer and supplemented with 400 µg of house-made TEV protease, and incubated overnight on a roller at 4 °C. The resulting flow-through was collected and concentrated using a pre-equilibrated 100K Amicon Ultra-15 concentrator (Millipore), then loaded onto a TSKgel G4000SWXL 7.8/300 column equilibrated with GF150 buffer (25 mM HEPES pH 7.2, 150 mM KCl, 1 mM MgCl2, 5 mM DTT, 0.1 mM ATP) at 4 °C. Peak fractions were pooled and concentrated to 2.5-3 mg/ml for immediate use in cryo-EM grid preparation. The quality of dynein is monitored using negative staining EM (**Extended Data Fig. 1**).

### α-registry human dynein motor domain purification

The human dynein motor domain locked in α-registry was expressed and purified from insect cells (ExpiSf9 cells; Life Technologies) as previously described(23). Briefly, 4 ml of ExpiSf9 cells at 2.5 × 106 cells/ml, which were maintained in ExpiSf CD Medium (Life Technologies), were transfected with 5-9 µg of bacmid DNA using ExpiFectamine (Gibco) according to the manufacturer’s instructions. 4-8 days following transfection, the cells were pelleted, and 1-2 ml of the resulting supernatant (P1) was used to infect 200 ml of ExpiSf9 cells (5 × 106 cells/ml). Approximately 65 hours later, the cells were harvested (2000 x g, 20 min), washed with lysis buffer (50mM HEPES, pH 7.4, 100 mM NaCl, 10% glycerol, 1mM DTT, 0.1mM Mg-ATP, 1mM PMSF), pelleted again (1810 x g, 20 min), and resuspended in an equal volume of same. The resulting cell suspension was drop frozen in liquid nitrogen and stored at −80ºC. For protein purification, 30 ml of additional lysis buffer supplemented with cOmplete protease inhibitor cocktail (Roche) was added to the frozen cell pellet, which was then rapidly thawed in a 37ºC water bath prior to incubation on ice. Cells were lysed in a dounce-type tissue grinder (Wheaton) using 50-60 strokes. Subsequent to clarification at 310,000 x g for 1 hour, the supernatant was applied to 2 ml of IgG sepharose fast flow resin (GE) pre-equilibrated in lysis buffer, and incubated at 4ºC for 3-5 hours. Beads were then washed 3 times with with 5-10 ml of lysis buffer, and 2 times with 5-10 ml of TEV buffer (50 mM Tris pH 7.4, 150 mM potassium acetate, 2 mM magnesium acetate, 1 mM EGTA, 10% glycerol, 1 mM DTT, 0.1 mM Mg-ATP). The beads were incubated with TEV protease overnight at 4ºC. The next morning, the recovered supernatant was collected, concentrated and used directly without freezing.

### Purification of yeast dynein motors

Purification of yeast dynein (wild-type and mutants; ZZ-2xTEV-6His-GFP-3HA-GST-dyneinMOTOR-HALO, all under the control of the galactose-inducible promoter, GAL1p) was performed as previously described(19, 91). Briefly, yeast strains were generated with plasmids encoding wild-type or mutant dyneins integrated into the yeast genome (after digestion with ApaI, as previously described(19, 91). The strains were grown in YPA supplemented with 2% galactose, harvested, washed with cold water, and then resuspended in a small volume of water. The resuspended cell pellet was drop frozen into liquid nitrogen and then lysed in a coffee grinder (Hamilton Beach). After lysis, 0.25 volume of 4X lysis buffer (1X buffer: 30 mM HEPES, pH 7.2, 50 mM potassium acetate, 2 mM magnesium acetate, 0.2 mM EGTA) supplemented with 1 mM DTT, 0.1 mM Mg-ATP, 0.5 mM Pefabloc SC (concentrations for 1X buffer) was added, and the lysate was clarified at 310,000 x g for 1 hour. The supernatant was then bound to IgG sepharose 6 fast flow resin (Cytinva) for 2-4 hours at 4°C, which was subsequently washed three times in 5 ml lysis buffer, and twice in TEV buffer (50 mM Tris, pH 8.0, 150 mM potassium acetate, 2 mM magnesium acetate, 1 mM EGTA, 0.005% Triton X-100, 10% glycerol, 1 mM DTT, 0.1 mM Mg-ATP, 0.5 mM Pefabloc SC). To fluorescently label the proteins, the bead-bound protein was incubated with either 10 µM JFX646-HaloTag or JFX549-HaloTag for 30-60 minutes at 4°C. The resin was then washed four more times in TEV digest buffer, then incubated in TEV buffer supplemented with TEV protease overnight at 4°C. Following TEV digest, the protein-containing supernatant was collected using a spin filtration device, aliquoted, flash frozen in liquid nitrogen, and stored at −80ºC.

### Microtubules Preparation

Porcine tubulin, obtained either from Cytoskeleton or purified in-house, was resuspended in MT buffer (25mM MES, 70mM NaCl, 1mM MgCl2, 1mM EGTA, and 1mM DTT, pH 6.5) to a final concentration of 10mg/ml, flash-frozen, and stored at −80°C. Microtubules were polymerized using a previously described protocol(8). In brief, tubulin was diluted in MT buffer supplemented with 3 mM GTP. The mixture was incubated on ice for 5 minutes and subsequently transferred to a 37°C incubator for 1 hour. Following polymerization, 5 µM paclitaxel (MilliporeSigma) was added to the mixture. The microtubules were then pelleted (20,000 rcf for 8 minutes at room temperature) and resuspended in MT buffer containing 5 µM paclitaxel.

### ATPase measurements

Basal and microtubule-stimulated ATPase activities were measured using the EnzChek phosphate assay kit (Invitrogen). Assays were performed in motility buffer (30 mM HEPES, pH 7.2, 50 mM potassium acetate, 2 mM magnesium acetate, 1 mM EGTA, 10% glycerol) supplemented with 1 mM Mg ATP, with or without 2 µM taxol-stabilized microtubules and 5 nM 6His-GST-dyneinMOTOR (wild-type or mutants). Reactions were initiated with the addition of dynein, and the absorbance at 360 nm was monitored by plate reader (BMG LabTech CLARIOstar Plus) for at least 20min. Background phosphate release levels (presumably from microtubules) for each reaction were measured for 2-5 min before addition of dynein motors. Relative ATPase levels were determined by fitting linear regressions to the time-resolved absorbance values, and determining the slope from each. Data shown are from 4-6 independent replicates, from two independent protein preparations.

### Single-molecule motility assays

Single-molecule motility assays were performed as previously described(19, 25) with minor modifications. In brief, flow chambers constructed using slides and plasma cleaned and silanized coverslips attached with double-sided adhesive tape were coated with anti-tubulin antibodies (8 µg ml1, YL1/2; Accurate Chemical Scientific Corporation) and then blocked with 1% Pluronic F-127 (Fisher Scientific). Taxol-stabilized microtubules assembled from unlabelled porcine tubulin (Cytoskeleton) were introduced into the chamber. After incubation for 2-5 min, the chamber was washed with motility buffer (see above) supplemented with 20 µM taxol and 1 mM DTT. Dynein, diluted in an oxygen-scavenging motility buffer (30 mM HEPES, pH 7.2, 50 mM potassium acetate, 2 mM magnesium acetate, 1 mM EGTA, 10% glycerol, 50 nM protocatechuate 3,4-dioxygenase, 2.5 mM protocatechuic acid, 1 mM Trolox, 1 mM cyclooctatetraene, 1 mM 4-nitrobenzyoyl alcohol) supplemented with 1 mM DTT, 20 µM taxol, and 1 mM Mg-ATP was then added. TIRFM images were immediately collected using an iLAS2 RING TIRF system on a Nikon Ti-2E inverted microscope equipped with a 1.49 NA 100X TIRF objective, a Ti2-SS-E motorized stage, piezo Z-control, and an iXon LIFE 897 cooled EM-CCD camera (Andor). 561 or 640 nm lasers housed in an LUN-F 3 (Nikon) were used along with a multi-pass quad filter cube set (C-TIRF for 405/488/561/638 nm; Chroma) and emission filters mounted in a filter wheel (600/50 nm and 700/75 nm; Chroma). To determine microtubule polarity (due to the diffusive/bidirectional behavior of the Y1902A mutant), wild-type GST-dyneinMOTOR-HaloJF549 (diluted in oxygen-scavenging motility buffer) was introduced into the imaging chamber immediately after acquiring a movie of the mutant GST-dyneinMOTOR-HaloJF646. To image non-fluorescent microtubules, we used interference reflection microscopy, as previously described (92). Images of dynein motors were acquired at 2 second intervals for 8 min. Velocity and run length values were determined from kymographs generated using the MultipleKymograph plugin for FIJI/ImageJ. Data shown are from 3 independent replicates, from 2 independent protein preparations.

### Cryo-EM sample and grid preparation for dynein unbound from microtubules

For dynein samples in AMPPNP, ATP-Vi, and ADP conditions, the concentrated dynein obtained after gel filtration was diluted with GF150 buffer containing 0.1 mM ATP to a concentration of 2.0 mg/ml. Subsequently, ATP-vanadate, AMPPNP, or ADP was added to a final concentration of 5 mM, and the mixture was incubated with dynein for 1 hour on ice before vitrification. In the case of dynein in ATP condition, the protein was incubated with 5 mM ATP on ice and immediately vitrified. For dynein in apo buffer (GF150 buffer without ATP), the protein was diluted into the apo buffer and concentrated using an Amicon concentrator. This concentration process was repeated three times, resulting in an estimated 8000-fold dilution of the original 0.1 mM ATP. The final concentration of dynein in the apo buffer was adjusted to 2 mg/ml for vitrification. Prepared samples were applied to Quantifoil holy carbon grids (R2/1, 300 mesh gold or R2/2, 200 mesh) and incubated in the chamber of the Vitrobot Mark IV unit (FEI) for 5 seconds at 4°C and 100% humidity. Subsequently, the grids were blotted for 3-5 seconds and plunged into liquid ethane.

### Cryo-EM sample and grid preparation for dynein bound to microtubules

For the reconstitution of dynein-MT complexes (**Extended Data Fig.2**) in apo condition, dynein in apo buffer was mixed with microtubules (MTs) at room temperature in MT-binding buffer (30 mM HEPES pH 7.2, 60 mM KCl, 1 mM EGTA, 5 mM MgSO4, 1 mM DTT, 5 µM paclitaxel). The dynein-MT complex was then pelleted at 20,000 rcf for 8minutes. The optimal resuspension volume, which determined the final concentration used for cryo-EM vitrification, was estimated using negative staining electron microscopy. For dynein-MT complexes in ADP or AMPPNP condition, the final MT binding buffer was supplemented with 5 mM ADP or 5 mM AMPPNP. A volume of 3.5 µL of the dynein-MT complex was applied to Quantifoil holy carbon grids (R2/1, 300 mesh gold), followed by a 4-second wait time, blotting for 3-6seconds, and subsequent freezing in liquid ethane using a Vitrobot Mark IV unit (FEI). The Vitrobot chamber was maintained at close to 95% humidity and 22°C during the process.

### Cryo-EM data collection

All cryo-EM grids underwent initial screening at the Yale ScienceHill-Cryo-EM facility using a Glacios microscope (Thermo Fisher Scientific). Subsequent cryo-EM data were collected at three different sites: Yale ScienceHill-Cryo-EM facility, utilizing a Glacios microscope operated at 300 keV with a K3 detector; Yale WestCampus-Cryo-EM facility, where a Krios microscope operated at 300 keV equipped with a K3 detector and a Bioquantum Energy Filter was utilized; and the Laboratory for BioMolecular Structure at BNL with a Krios microscope operated at 300 keV with a K3 detector and a Bioquantum Energy Filter. Automatic data collection was facilitated by either SerialEM(93) or EPU software. In total, 33,302, 6,016, 7,488, 8,355, 5,920, and 5,884 movies were acquired for dynein-ATP, dynein-ATPVi, dynein-AMPPNP-lowMg^2+^, dynein-AMPPNP-high Mg^2+^, dynein-ADP, and dynein-Apo off microtubules, respectively. Additionally, 9,935, 11,828, and 5,690 movies were acquired for dynein-Apo-MT, dynein-ADP-MT, and dynein-AMPPNP-MT on microtubules, respectively. Detailed data collection parameters can be found in supplementary tables S1-S7 corresponding to different datasets.

### Cryo-EM image processing for dynein unbound from microtubules

Preprocessing steps, including motion correction, CTF estimation, and particle picking, were conducted either in cryoSPARC Live(94) or via an in-house script utilizing MotionCor2(95), GCTF(96), and Gautomatch. Cryo-EM scripts for real-time data transfer and on-the-fly preprocessing are available for download at https://github.com/JackZhang-Lab.

For the dynein-ATP off microtubules dataset (**Extended Data Fig. 1, 2**), the blob picker was utilized to select motor domains in phi or open conformations, with the cross-correlation score set to 0.01 to enable sampling of all possible motors during the picking step. A total of 8,534,500 particles were extracted using a box size of 400 pixels, subsequently binned to 200 pixels (1.664 Å pixel size) for faster computation. From this pool, a subset of 640,000 particles underwent subsequent rounds of 2D classification, ultimately yielding 164,658 high-quality particles selected for ab initio reconstruction (6 classes). During this phase, phi motor domains and several single motor domains were identified. Following the initial model acquisition, all original particles were employed for subsequent heterogeneous refinement to prevent the loss of motor particles during 2D classification. The original particles were divided into four subsets, with each subset subjected to heterogeneous refinement (12 classes). This initial heterogeneous refinement process was iterated three times, with updated references provided each time, successfully identifying phi-motors, single motor domains with bent or straight linkers, and several junk classes.

Each motor class from four subsets was merged for additional heterogeneous refinement or 3D classification without alignment. This step further explores the dynamics of the motor domain. For example, state-4, characterized by a straight linker, was initially obscured within the state-2 class due to their closed motor ring conformation. Similarly, state-5, state-6, and state-7 were initially grouped into a single class and later separated during subsequent heterogeneous refinement. The second-level heterogeneous refinement was conducted three times, with updated references provided each time. Further classification of individual classes ceased to reveal novel conformations due to limited particle numbers and achievable resolution. An increase in dataset size may unveil additional intermediate or transient conformations.

Following homogeneous refinement, most classes achieved resolutions at the Nyquist limit of 3.3 Å. Subsequently, particles were re-extracted at the full box size of 400 pixels (0.832 Å). For high-resolution refinement, two rounds of global and local CTF refinement followed by local refinement were performed. The resolution of single motor domains from open dynein reached 2.9-3.6 Å, while phi-motor domains achieved 2.6 Å using C2 symmetry. To further enhance the resolution of phi-motor domains, symmetry expansion of the phi-motor domain was executed to account for inter-motor domain dynamics. Particles were recentered on one of the motor domains and re-extracted using a box size of 640 pixels to incorporate delocalized high-resolution information. After localized refinement and another round of global and local CTF refinement, the phi-motor domain achieved 2.25 Å resolution.

For full-length phi dynein reconstruction, two sets of particles were considered. Firstly, junk particles from the initial refinement were merged and subjected to heterogeneous refinement using the dynein tail map (EMD-3703(15)) and junk volumes as references, yielding classes with recognizable tail domains. Secondly, particles from phi dynein motor domains were recentered at the tail and re-extracted (box size 400 pixels binned to 200 pixels). These two sets of particles were merged for additional heterogeneous refinement focusing on the tail region, resulting in classes with clear tail domain features. The selected class was re-extracted using a large box size of 1024 pixels (binned to 256 pixels) for full-length dynein refinement. Duplicated particles were removed during re-extraction. After another heterogeneous refinement, the class with both stable tail and motor domain was selected and refined to 6.8 Å resolution. To improve the resolution of the tail domain, particles were recentered on the tail and re-extracted using a box size of 384 pixels. A mask covering only the tail domain was created, and after local refinement, the entire tail domain reached 4.3 Å resolution, albeit with large variations in resolution among different tail regions. Masks covering the NDD region, middle, and neck regions were used to improve local resolution. The middle region of the tail was refined to 3.8 Å, revealing side chain density, while the NDD and neck regions reached 6.5 Å with resolved secondary structures. Localized refined maps were fitted to the consensus map and stitched together in ChimeraX(97) using the “volume maximum” command to create a composite map of full-length human phi dynein. The resolutions of all maps were estimated by Fourier shell correlation calculations(98, 99) embedded in cryoSPARC. Local resolution analysis(100) was performed in cryoSPARC.

Datasets of dynein under ATP-Vi, AMPPNP, ADP, and apo conditions underwent processing using the same procedures. The simple flowchart and map quality are summarized in Extended Data Figs. 4, 5.

### Cryo-EM image processing of dynein bound to microtubules

For cryo-EM image processing of dynein-MT datasets, we employed a pipeline previously developed for MT tracing and MT-signal subtraction (**Extended Data Figs. 3, 5**)(73, 74). Initially, MT particles were detected using template matching with a distance cut-off of 8 nm, with a relatively low cross-correlation score set to include MT particles with low contrast or located at cross-over locations. Following three rounds of 2D classification, false-picked MT particles, including carbon edges, and mis-centered MTs were rejected. The metadata of the selected particles was converted to Relion style using “csparc2star.py” in pyem (https://zenodo.org/records/3576630), and the coordinates were recentered and subjected to “multi-curve fitting”. After MT-signal subtraction, original micrographs were replaced by MT-signal subtracted micrographs.

Subsequently, the blob-picker was used to detect motor particles in these new micrographs, which were then extracted using a box size covering the single motor domain. 2D classification was utilized to remove junk particles and residual MT particles, after which the selected particles underwent ab initio reconstruction. Similar to our above workflow, all original particles were used for initial heterogeneous refinement. 3D classification-based particle sorting greatly aided in identifying motor particles with low signal-to-noise ratios due to residual MT signal influence. After several rounds of sorting, classes with clear motor features were selected and subjected to local refinement, global, and local CTF refinement. Notably, the post-1 linker-docking mode was observed in three datasets, while the post-2 was exclusively found in the dynein-MT-ADP dataset. The resolution of individual motor domains ranged from 2.9 Å to 3.6 Å.

For the reconstruction of full-length dynein bound to MT, particles were reextracted using a large box size covering two motor domains. The particles were directly reconstructed into a volume featuring the weak density of the second motor. Subsequently, this map was low-pass filtered to 60 Å for heterogeneous refinement, where some classes exhibited enhanced density for the second motor. We then manually fitted the motor map into the weak density to create a two-motor map. This map, together with previous reconstructions, was used for another round of heterogeneous refinement. The newly generated maps served as new references for iterative rounds of heterogeneous refinement. Ultimately, three main states were predominantly observed for each dynein-MT dataset: i) two stable and parallel heads, ii) one stable leading head, and iii) one stable trailing head. However, due to the limited number of particles and resolution, we were unable to detect two motors with different step sizes in our dynein-ADP-MT datasets.

Additionally, we attempted to reconstruct dynein-MTBD-MT by reverting the MT signal using the original micrographs. Unlike our previously determined out-arm dynein bound to protofilament structures featuring defined tubulin subunit density(54), the protofilament of dynein-1 bound to MT exhibited smeared density. Furthermore, subsequent 3D classification without alignment failed to generate a relatively stable stalk-MTBD-protofilament class due to the limited number of particles. These limitations precluded us from performing localized refinement on the MTBD-MT region.

### Model building and refinement

For model building and refinement, we utilized two previously reported human dynein motor structures (PDB: 5NUG(15), 7Z8G(8)) as initial models for different states of dynein in ATP. Individual domains (linker, AAA large, AAA small) were rigidly docked into the cryo-EM map in UCSF ChimeraX. Subsequently, Namdinator(101), a molecular dynamics flexible fitting tool, was employed to further refine the model into the cryo-EM map. Manual inspection and adjustments of the model were carried out in COOT v0.9.5(102, 103).

For water model building in the phi-motor, the maps underwent sharpening using two methods: i) different B-factor sharpening with local filtering based on local resolution in cryoSPARC, and ii) density modification with phenix.resolve-cryo-em(104) using only maps as input. The processed maps were up-sampled with a pixel size of 0.416 Å and cropped to a box size of 400 pixels. Initial solvent water molecules were placed using phenix.douse, resulting in approximately 2358 automatically placed water molecules. Subsequent manual inspection in COOT led to the addition or deletion of water molecules, resulting in a final placement of 1897 water molecules.

All models underwent iterative refinement using Phenix real-space refinement 1.21rc1-5190 and manual rebuilding in COOT. These refined models were utilized as initial models for dynein in other nucleotide conditions or dynein bound to MTs. The quality of the refined models was assessed using MolProbity(105) integrated into Phenix(104), with statistics reported in Supplementary Table 1–8.

### Vector map analysis of dynein motor domains

For vector map in Figures 2d and 5b, we used the “PDBArrows” script from https://github.com/sami-chaaban/PDBArrows. In brief, the isolated AAA domains or entire motor domains were aligned in ChimeraX (two structures per vector map). After alignment, the two structures were saved into the same coordinate system. “PDBArrows” was used to generate sequences of the two structures for multiple sequence alignment using ClustalW (https://www.genome.jp/tools-bin/clustalw). Even though the two structures are both human dynein, this step is necessary since one structure may be missing some residues during model building. A ‘.bild’ file (a vector map of paired C-alpha carbon distances) was then generated with the script using the two PDBs and sequence alignment files as inputs. The .bild file was then opened in ChimeraX, which shows the final vector map, and the color command (for example “.color 0000ff”) was used to color the individual domains.

### Plots, and molecular graphics

Plots were generated using GraphPad Prism. Molecular graphics were prepared using UCSF ChimeraX(97).

---

**Extended Data Figure 1.**
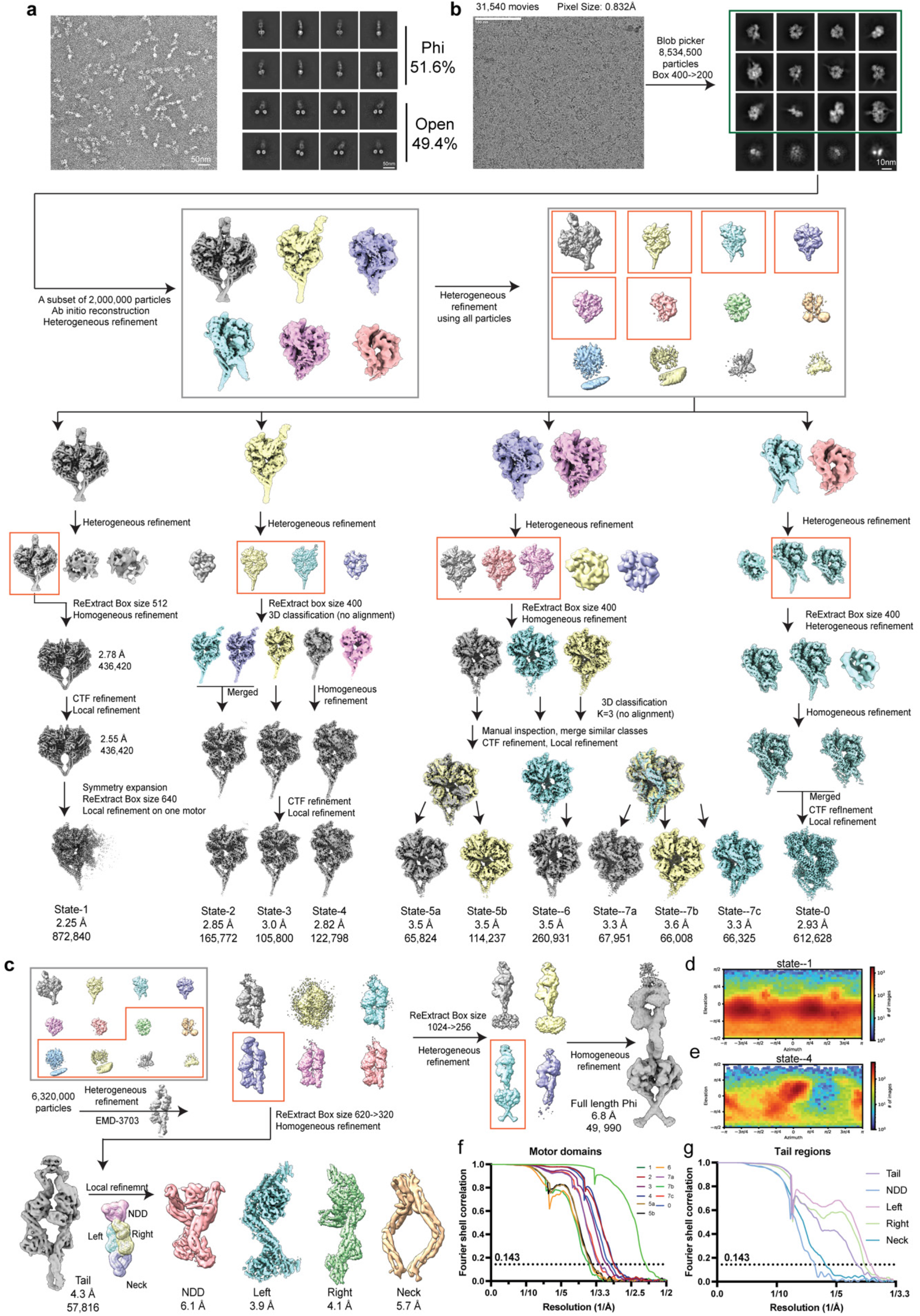
Cryo-EM data processing of full-length human dynein-1 in the presence of 5 mM ATP. **(a)** A representative negative staining micrograph of purified full-length human dynein-1 and corresponding 2D class averages. **(b)** A typical cryo-EM micrograph and the flowchart of cryo-EM image processing. **(c)** Image processing of full-length phi dynein and tail region. **(d, e)** Orientation distribution of phi dynein and a representative state from open dynein. **(f, g)** The Fourier shell correlation (FSC) curves of the motor domain in different states and the tail domain. Datasets for dynein in other nucleotide conditions (ATP-Vi, AMPPNP, ADP, apo) were processed similarly.

**Extended Data Figure 2.**
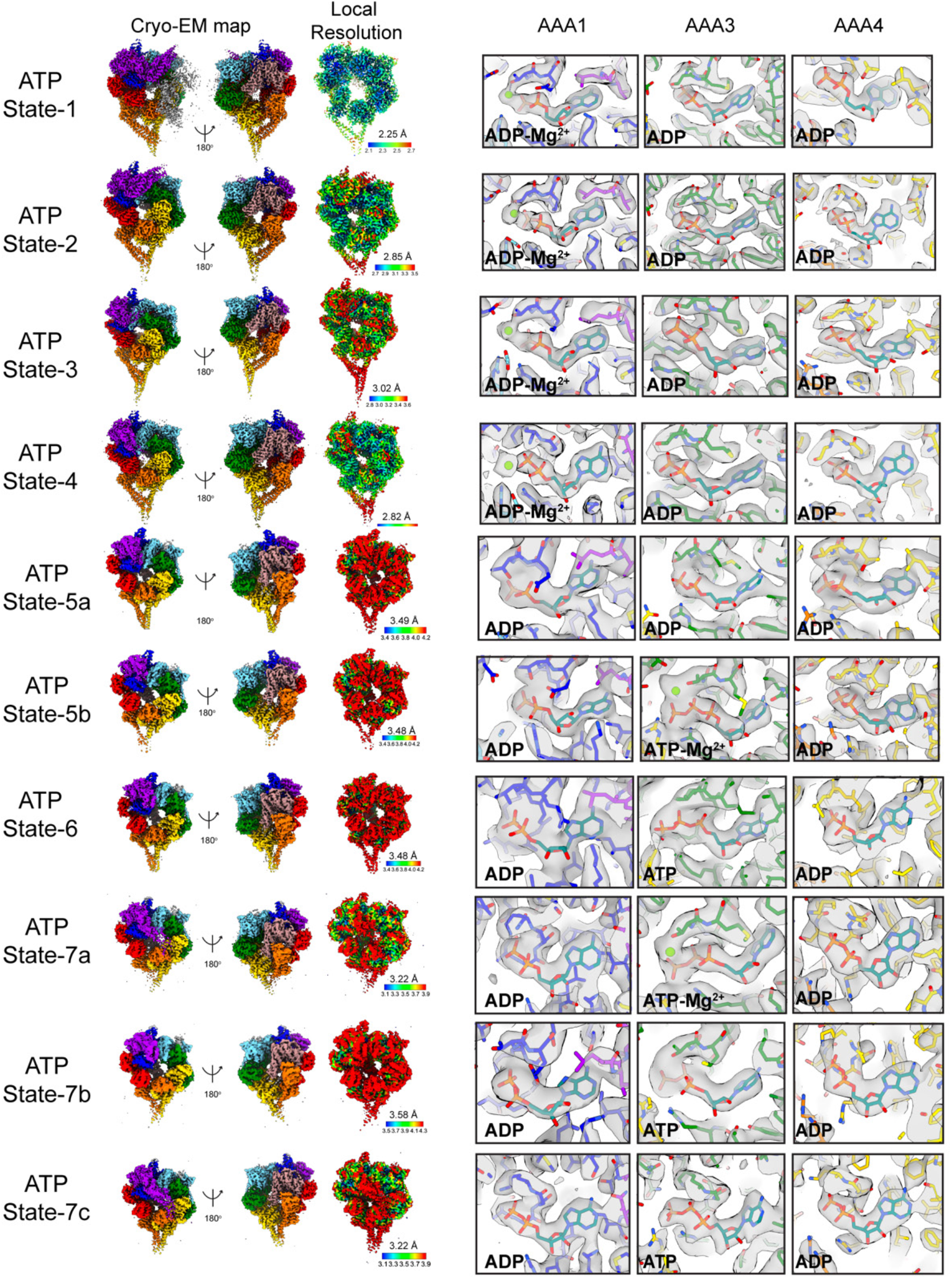
Local resolution analysis and identification of bound nucleotides in AAA1, AAA3, and AAA4 in different states from the dynein-ATP dataset. The color scheme for the motor domain is consistent with Figure 5.

**Extended Data Figure 3.**
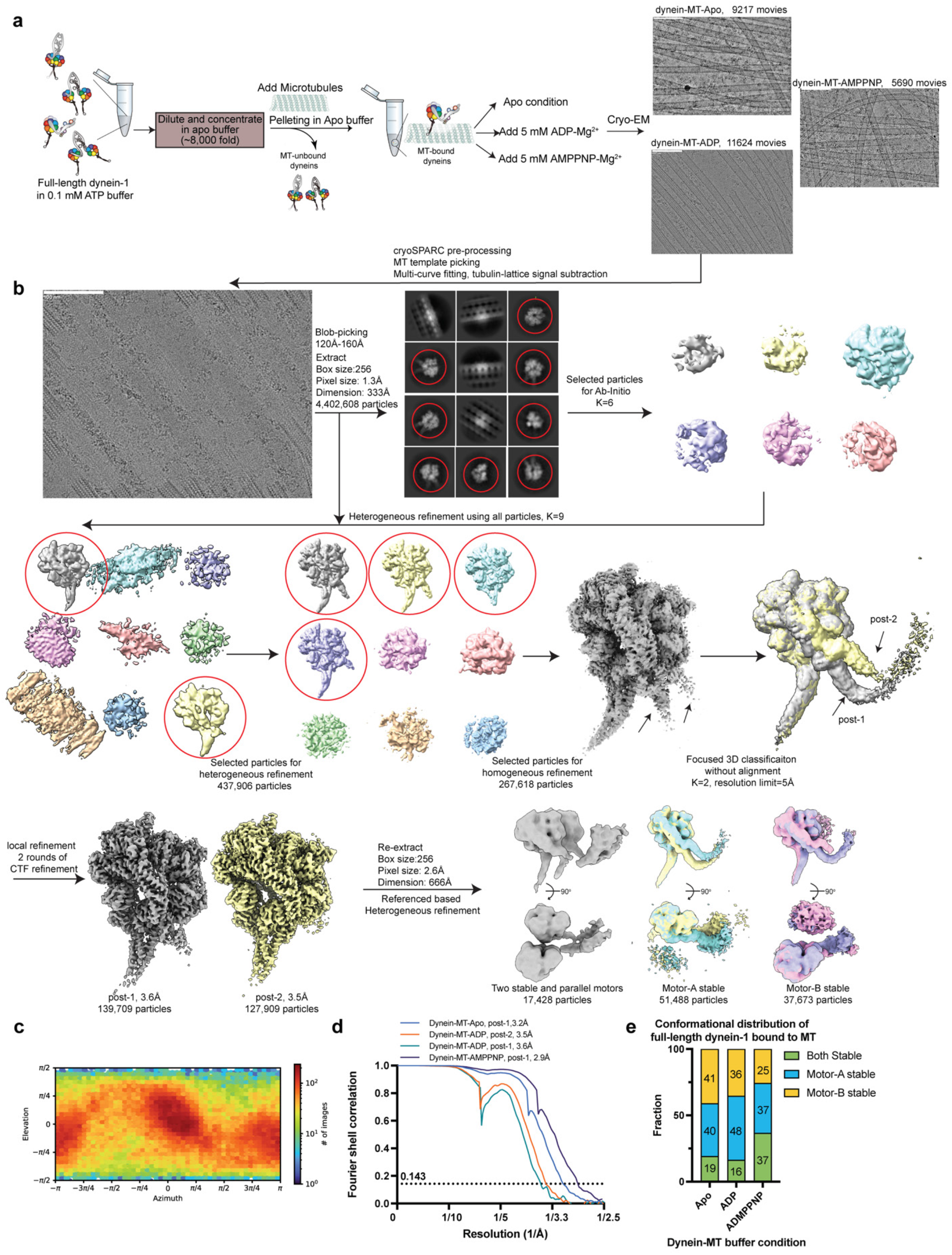
Cryo-EM data processing of full-length human dynein-1 bound to MTs. **(a)** Flowchart of sample preparation and typical cryo-EM micrographs of dynein bound to MTs in apo, ADP, and AMPPNP conditions. **(b)** Microtubule signal subtraction and image processing flowchart. **(c)** Orientation distribution of dynein-MT-ADP reconstruction. **(d)** FSC curves of four MT-bound motors. **(e)** Plot for the full-length dynein bound to MTs in different conformations (two stable heads, one stable trailing head, and one stable leading head).

**Extended Data Figure 4.**
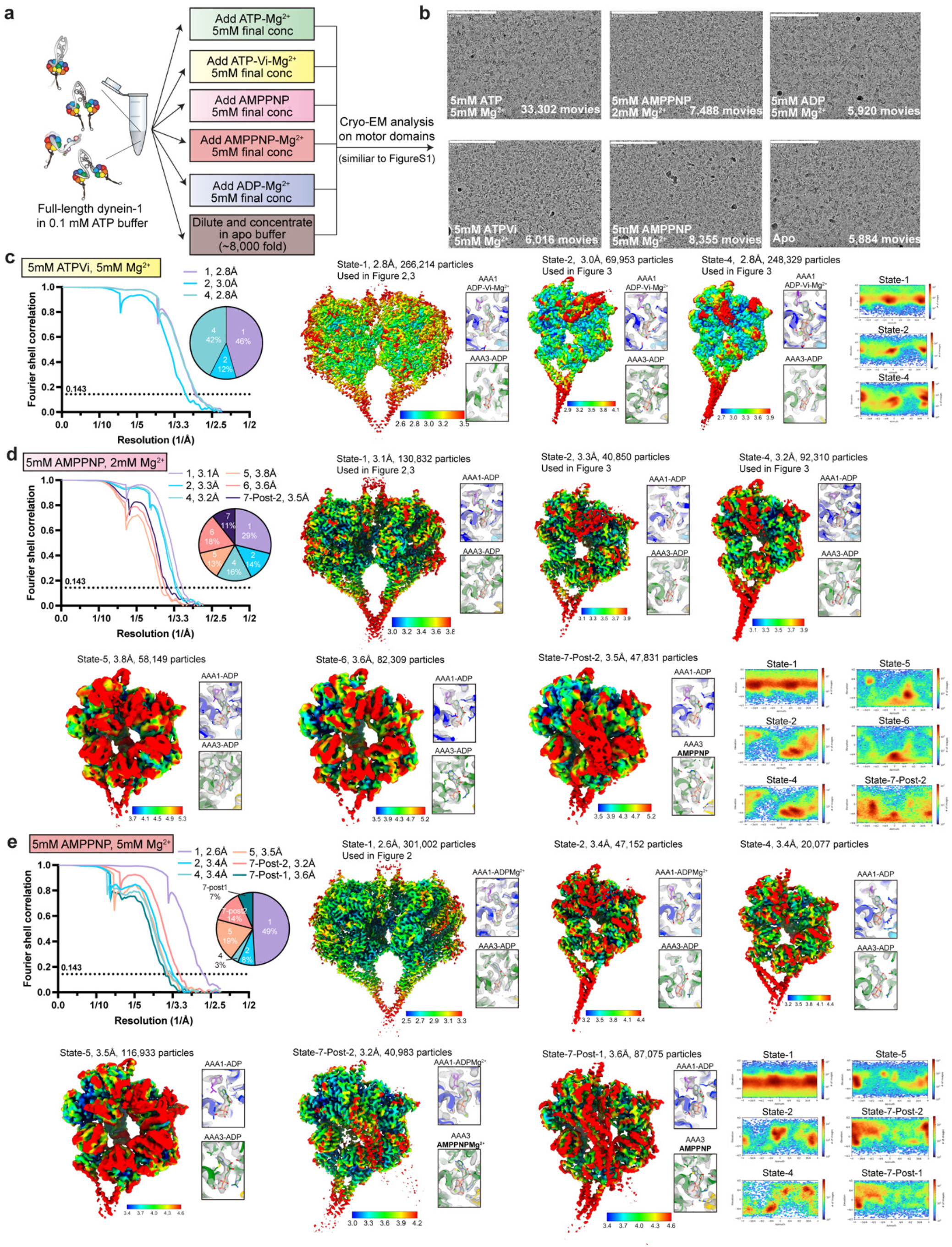
Cryo-EM analysis of dynein in ATP-Vi and AMPPNP conditions. **(a)** Cartoon schematic depicting the experimental method for cryo-EM analysis of full-length dynein in different nucleotide conditions. **(b)** Typical cryo-EM micrographs of dynein in different nucleotide conditions. **(c, d, e)** FSC curves and local resolution analysis of dynein in ATP-Vi, AMPPNP-low Mg^2+^, and AMPPNP-high Mg^2+^ conditions.

**Extended Data Figure 5.**
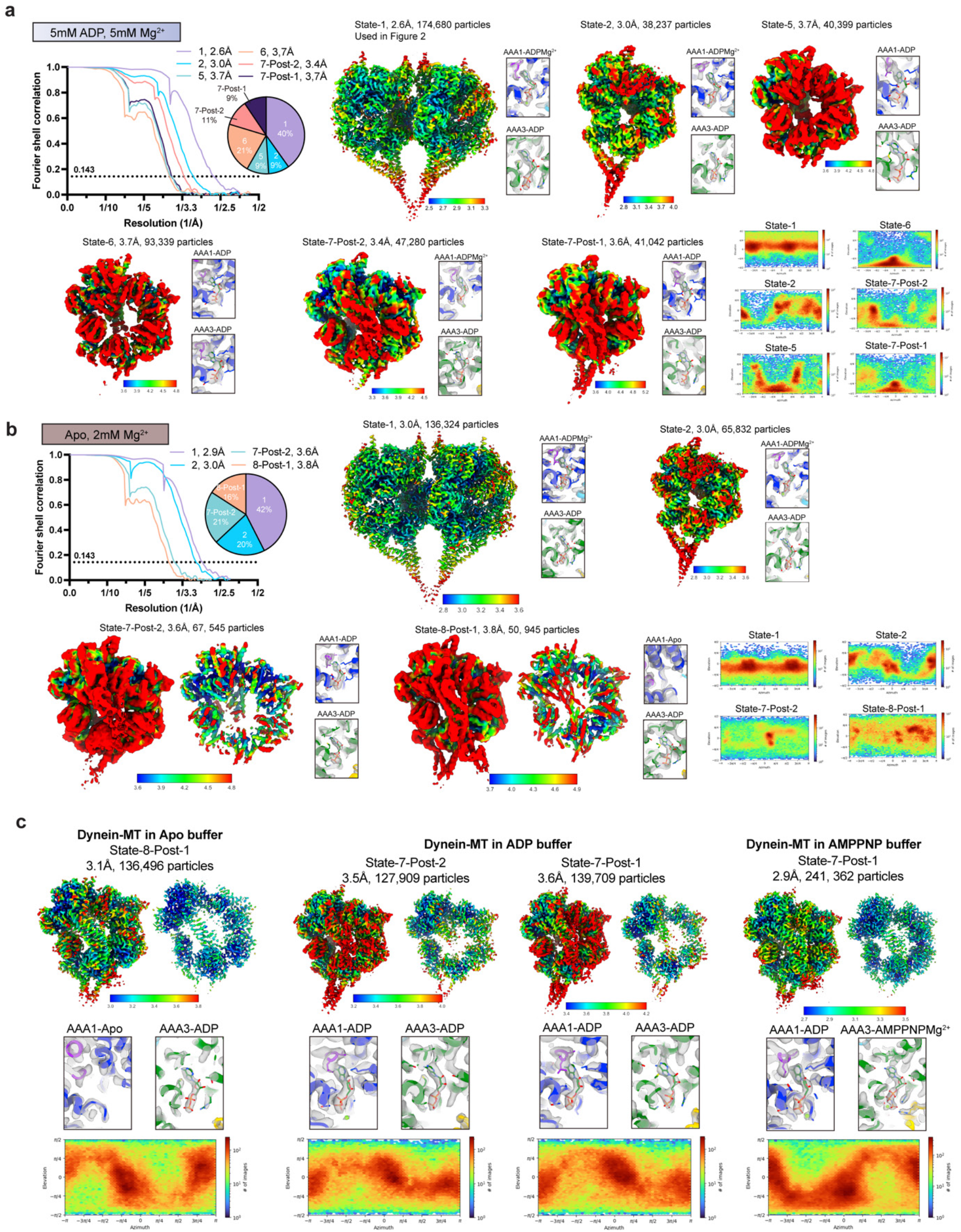
Cryo-EM analysis of dynein in ADP and apo conditions and dynein-bound to microtubules. **(a, b)** FSC curves and local resolution analysis of dynein in ADP and apo conditions. **(c)** FSC curves and local resolution analysis of dynein-bound to microtubules.

**Extended Data Figure 6.**
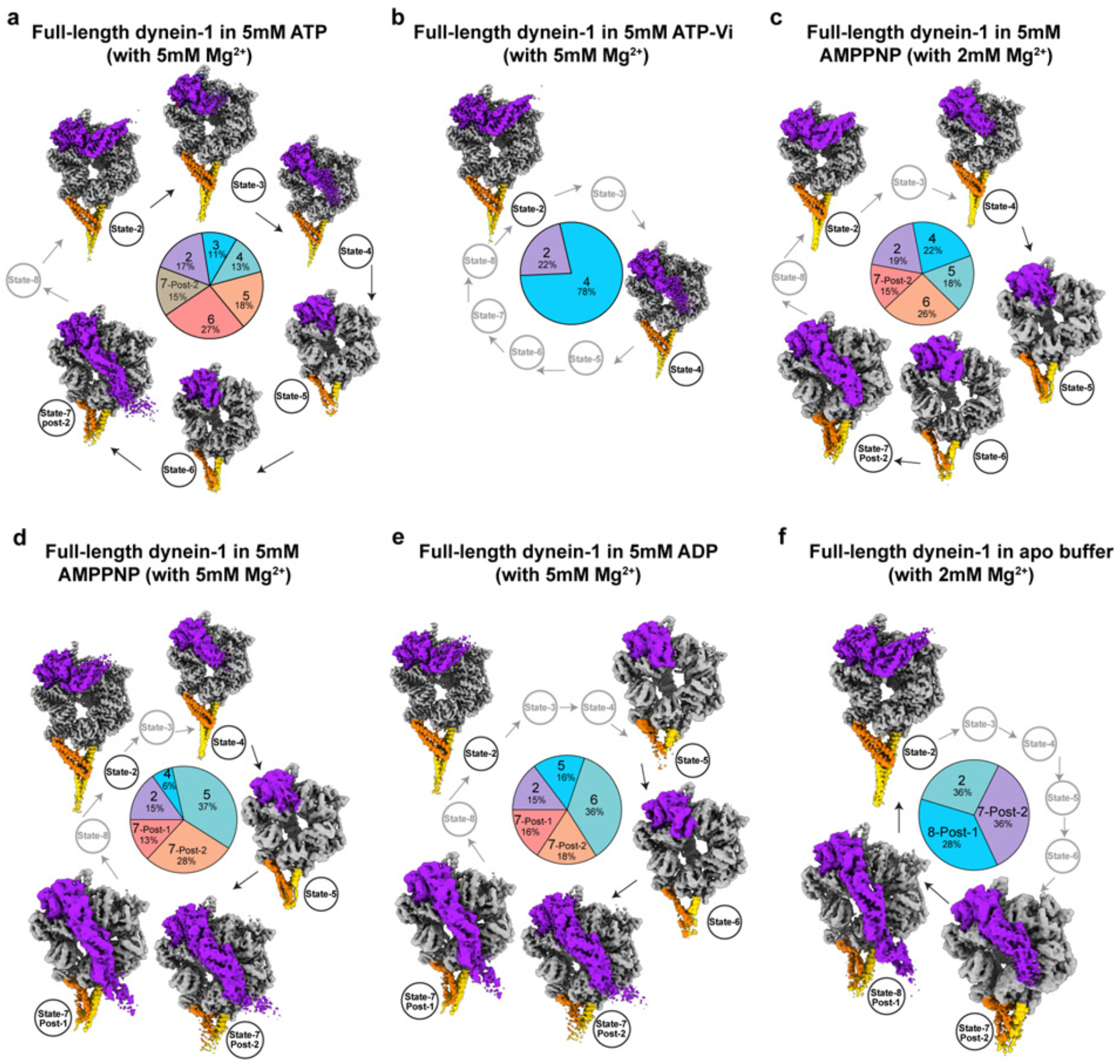
The conformational landscapes of active cycle motor domains in different nucleotide conditions. **(a-f)** Conformation landscapes of motor domains in ATP, ATP-Vi, AMPPNP, ADP, and apo conditions. Arrows indicate mechanochemical pathways. Percentages indicate the proportion of particles in each state. The missing states in each condition are shown in grey. Phi dynein is not included in this figure.

**Extended Data Figure 7.**
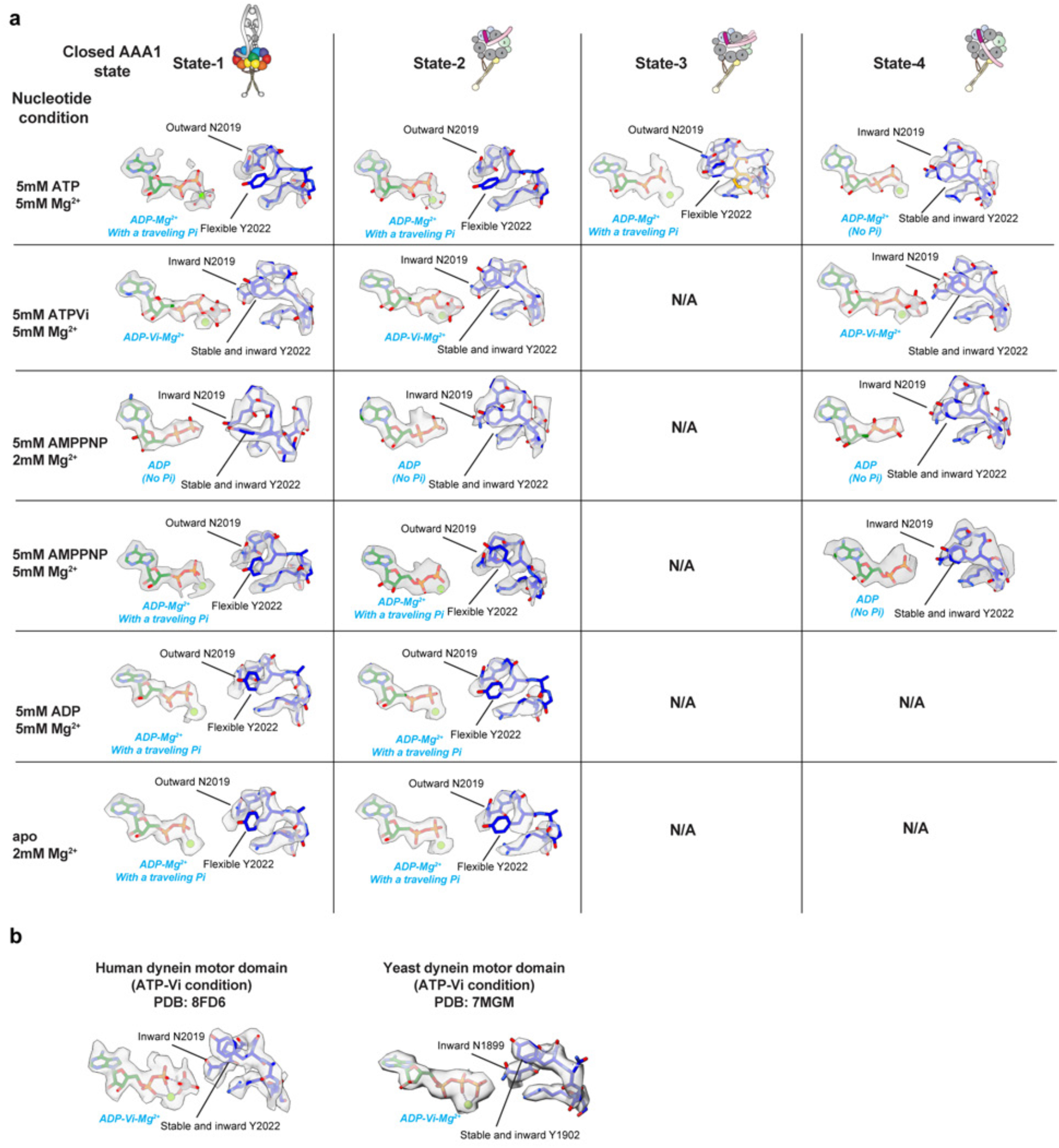
Cryo-EM densities of the nucleotide and sensor-I loop in various closed AAA1 states. **(a)** From this work. **(b)** From previously published structures (PDB:8FD6^21^, 7MGM^77^). The conformations of N2019 and Y2022 are highlighted.

**Extended Data Figure 8.**
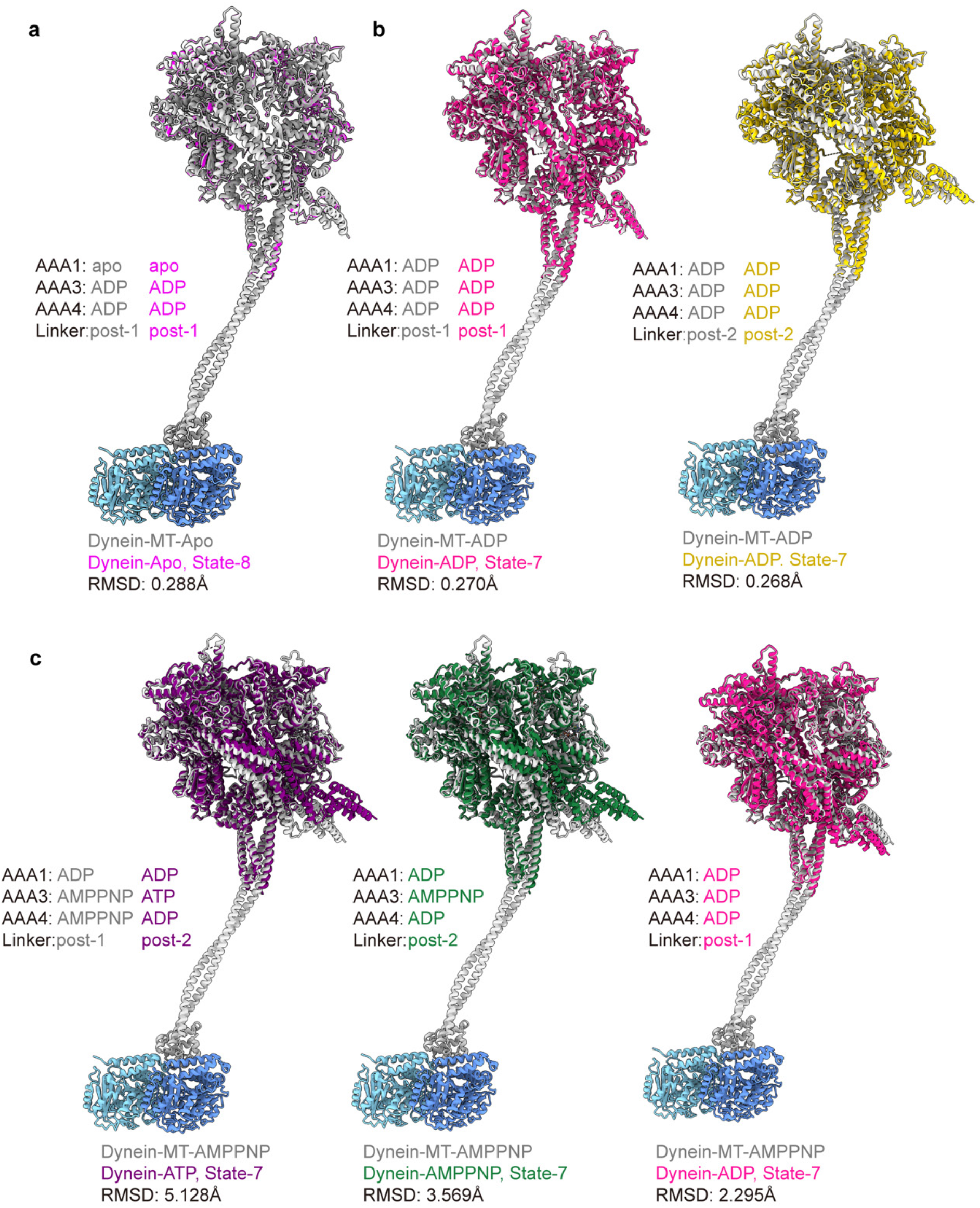
Structural comparisons of MT-bound and -unbound dynein motors. The nucleotide states in AAA1, AAA3, and AAA4 and the linker docking mode are listed. The MT-bound dynein motor domains (from linker to C-terminus) were used as references for structural fitting in ChimeraX.

**Extended Data Figure 9.**
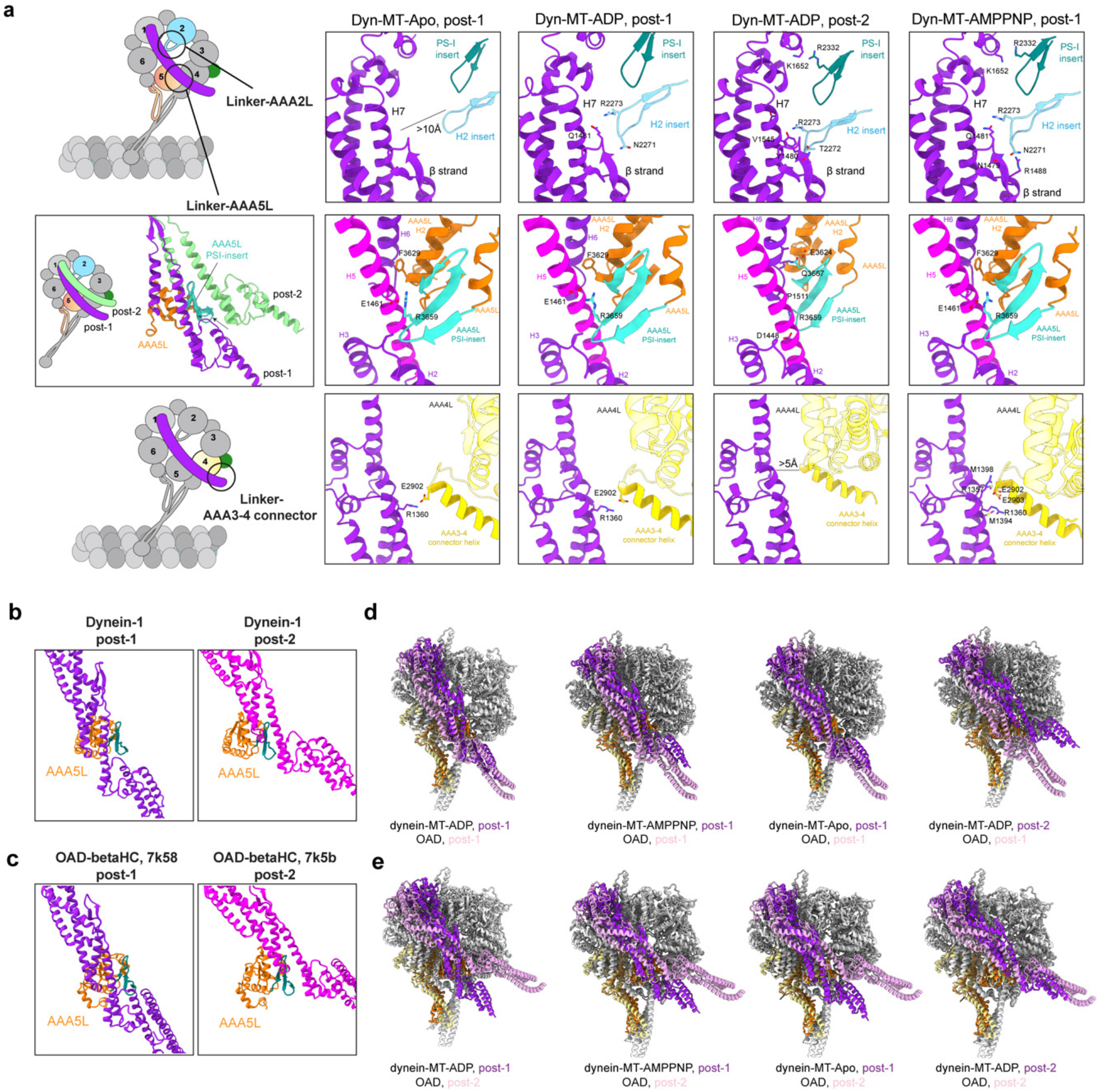
Systematic analysis of linker-ring interactions among four MT-bound motor domains. **(a)** Cartoon and molecular models showing the interaction interfaces including linker-AAA2L, linker-AAA5L, and linker-AAA3-4-connector helix. **(b-e)** Comparison of linker docking modes between dynein-1 and outer-arm dynein.

**Extended Data Figure 10.**
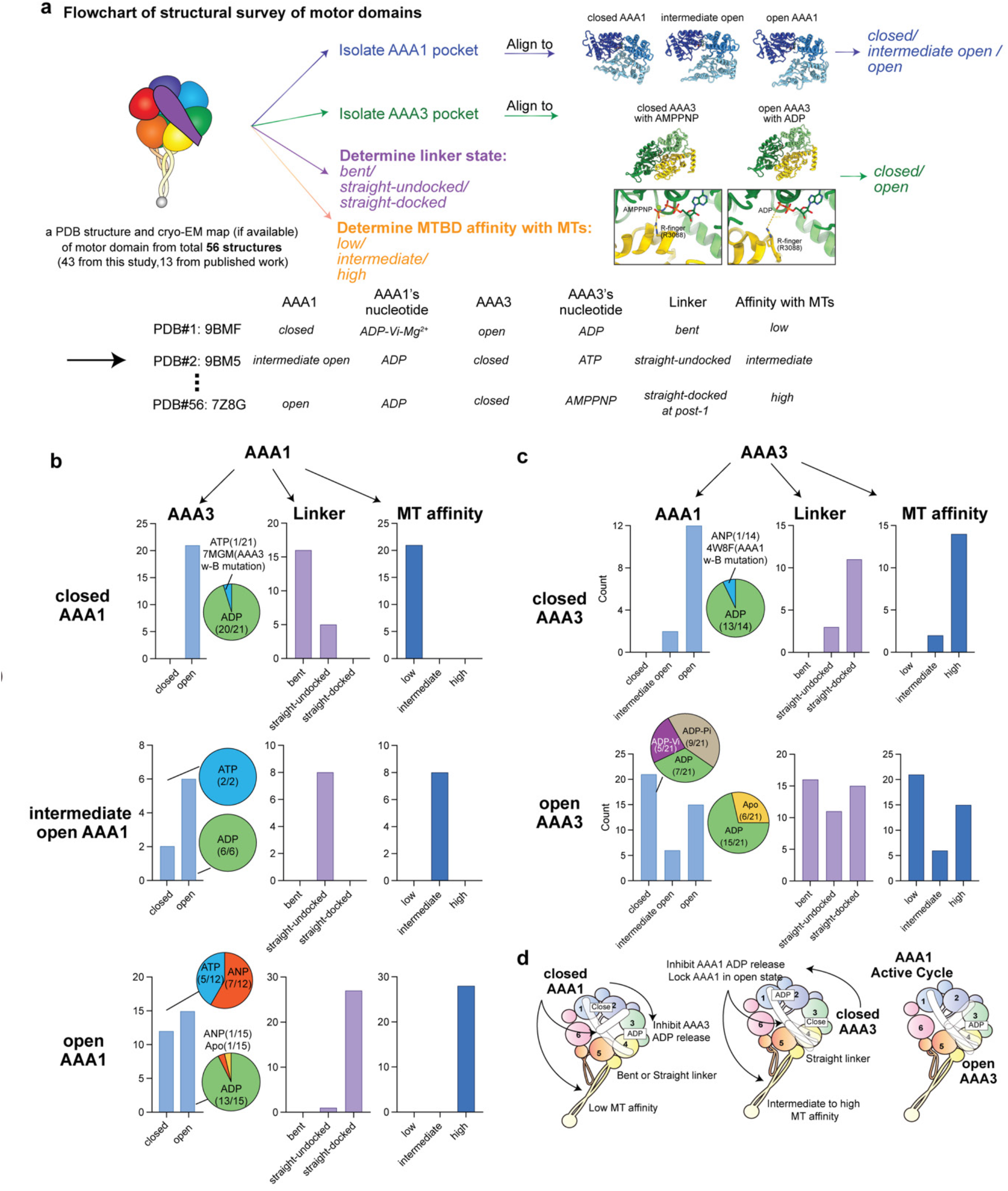
Structural survey of motor domains and statistical analysis of crosstalk between AAA1 and AAA3. **(a)** Flowchart of structural survey of 56 motor domain structures. **(b)** Statistical graphs showing how AAA1 communicates to AAA3, linker and MTBD. **(c)** Statistical graphs showing how AAA3 communicates to AAA1, linker and MTBD. **(d)** Cartoon model summarizing communication between AAA1, AAA3, linker and MTBD.

**Supplementary Video 1:** High-resolution cryo-EM map of phi dynein and representative local density maps.

**Supplementary Video 2:** High-resolution cryo-EM reconstruction of dynein bound to MTs.

**Supplementary Video 3:** The release of Pi from dynein AAA1 through the backdoor mechanism, triggering the linker straightening.

**Supplementary Video 4:** Communication between AAA1 pocket dynamics and MTBD.

**Supplementary Video 5:** The forward-stepping mechanism of dynein along MTs.

**Supplementary Table 1.** Cryo-EM data collection, refinement, and validation statistics of full-length dynein-1 in the presence of 5 mM ATP with 5 mM Mg^2+^.

|  |  |  |  |  |  |  |  |  |  |  |  |
| --- | --- | --- | --- | --- | --- | --- | --- | --- | --- | --- | --- |
| Description | Full-length dynein-1 in the presence of 5 mM ATP with 5 mM Mg<sup>2+</sup> |  |  |  |  |  |  |  |  |  |  |
|  | Composite map of full-length dynein-1 | State-1 Phi-motor | State-2 | State3 | State-4 | State-5a | State-5b | State-6 | State-7a-post-2 | State-7b-post-2 | State-7c-post-2 |
|  | EMD-44681 | EMD-44682 | EMD-44683 | EMD-44684 | EMD-44685 | EMD-44686 | EMD-44687 | EMD-44688 | EMD-44689 | EMD-44690 | EMD-44691 |
|  | PDB-9BLY | PDB-9BLZ | PDB-9BM0 | PDB-9BM1 | PDB-9BM2 | PDB-9BM3 | PDB-9BM4 | PDB-9BM5 | PDB-9BM6 | PDB-9BM7 | PDB-9BM8 |
| Data Collection and Processing |  |  |  |  |  |  |  |  |  |  |  |
| Facility | Yale West Campus-Cryo-EM facility |  |  |  |  |  |  |  |  |  |  |
| Microscope | Titan Krios |  |  |  |  |  |  |  |  |  |  |
| Voltage (kV) | 300 |  |  |  |  |  |  |  |  |  |  |
| Camera | K3 |  |  |  |  |  |  |  |  |  |  |
| Magnification | 105k |  |  |  |  |  |  |  |  |  |  |
| Pixel Size (Å) | 0.416 (super resolution) |  |  |  |  |  |  |  |  |  |  |
| Total Electron Exposure (e-/Å²) | 40 |  |  |  |  |  |  |  |  |  |  |
| Defocus Range (µm) | 1.5-2.7 |  |  |  |  |  |  |  |  |  |  |
| Symmetry Imposed | C1 | C1* | C1 | C1 | C1 | C1 | C1 | C1 | C1 | C1 | C1 |
| Num of mics | 33302 |  |  |  |  |  |  |  |  |  |  |
| Initial Particles | 4 670 501 |  |  |  |  |  |  |  |  |  |  |
| Final Particles | 57,816 | 872,840 | 165,772 | 105,800 | 122,798 | 65,824 | 114,327 | 260,931 | 67,951 | 66,008 | 66,325 |
| Refinement |  |  |  |  |  |  |  |  |  |  |  |
| Initial models | 5NUG, 7Z8G ab initio | 5NUG ab initio | 5NUG ab initio | 5NUG ab initio | 5NUG ab initio | 5NUG ab initio | 5NUG ab initio | 5NUG ab initio | 5NUG ab initio | 5NUG ab initio | 5NUG ab initio |
| Map pixel size | 1.664 | 0.416 | 0.832 | 0.832 | 0.832 | 0.832 | 0.832 | 0.832 | 0.832 | 0.832 | 0.832 |
| Map Resolution (Å) (FSC 0.143) | 3.3 | 2.25 | 2.85 | 3.02 | 2.82 | 3.49 | 3.48 | 3.48 | 3.22 | 3.58 | 3.22 |
| Map sharpening B-factor (Å²) | 0 | 60 | 76.2 | 56.4 | 72.8 | 63.9 | 63.8 | 56.5 | 62.9 | 59.9 | 63.3 |
| Model Composition |  |  |  |  |  |  |  |  |  |  |  |
| Non-hydrogen atoms | 89407 | 25271 | 23712 | 23061 | 23048 | 21869 | 21803 | 21782 | 23123 | 21829 | 23121 |
| Protein residues | 11063 | 2899 | 2,937 | 2,859 | 2,859 | 2,711 | 2,701 | 2,698 | 2,859 | 2,704 | 2,859 |
| Ligands | MG:4<br>ADP:6<br>ATP:2 | MG:2<br>ADP:3<br>ATP:1<br>Water: 1897 | MG:2<br>ADP:3<br>ATP:1<br>Water:3 | MG:2<br>ADP:3<br>ATP:1 | MG:2<br>ADP:3<br>ATP:1<br>Water:7 | ADP:3<br>ATP:1 | MG:1<br>ADP:2<br>ATP:2 | ADP:2<br>ATP:2 | MG:1<br>ADP:2<br>ATP:2 | ADP:2<br>ATP:2 | ADP:2<br>ATP:2 |
| Model vs. Data |  |  |  |  |  |  |  |  |  |  |  |
| FSC Map to Model (Å) (FSC 0.5) | 3 | 2.4 | 2.8 | 3.1 | 2.8 | 3.8 | 3.8 | 4.0 | 3.5 | 4.0 | 3.5 |
| Correlation coefficient (mask) | 0.7 | 0.86 | 0.85 | 0.90 | 0.87 | 0.82 | 0.82 | 0.76 | 0.86 | 0.85 | 0.85 |
| **B factors (Å²)** |  |  |  |  |  |  |  |  |  |  |  |
| Protein | 100.96 | 25.33 | 39.98 | 50.59 | 33.75 | 89.22 | 101.82 | 85.68 | 53.63 | 114.60 | 44.50 |
| Ligand | 1.53 | 6.87 | 12.02 | 21.32 | 10.08 | 64.61 | 68.47 | 60.08 | 40.73 | 87.63 | 30.44 |
| Water | / | 21.56 | 11.75 | / | 11.54 | / | / | / | / | / | / |
| **R.m.s deviation** |  |  |  |  |  |  |  |  |  |  |  |
| Bond length (Å) | 0.006 | 0.004 | 0.002 | 0.002 | 0.002 | 0.004 | 0.003 | 0.005 | 0.003 | 0.004 | 0.003 |
| Bond angles (°) | 1.083 | 0.662 | 0.491 | 0.489 | 0.51 | 0.646 | 0.635 | 0.864 | 0.568 | 0.656 | 0.583 |
| **Validation** |  |  |  |  |  |  |  |  |  |  |  |
| Molprobity score | 1.94 | 1.51 | 1.4 | 1.38 | 1.39 | 1.64 | 1.7 | 1.91 | 1.63 | 1.80 | 1.62 |
| Clashscore | 13.53 | 9.59 | 7.3 | 6.87 | 7.05 | 12.42 | 12.9 | 21.04 | 9.3 | 13.52 | 9.43 |
| Rotamer outliers (%) | 0.24 | 0.04 | 0.84 | 0.00 | 0.04 | 0.17 | 0.54 | 0.13 | 0.31 | 0.54 | 0.00 |
| **Ramachandran plot** |  |  |  |  |  |  |  |  |  |  |  |
| Outliers (%) | 0.39 | 0 | 0 | 0 | 0 | 0 | 0.04 | 0 | 0.11 | 0.07 | 0.04 |
| Allowed (%) | 3.91 | 1.49 | 1.19 | 1.61 | 1.3 | 2.15 | 2.31 | 2.42 | 2.64 | 2.79 | 2.60 |
| Favored (%) | 95.7 | 98.51 | 98.81 | 98.39 | 98.70 | 97.85 | 97.66 | 97.58 | 97.26 | 97.14 | 97.36 |
| **Rama-Z (whole)** | 0.55 | 1.4 | 1.69 | 2.17 | 2.24 | 1.17 | 1.44 | 0.99 | 1.36 | 1.71 | 1.71 |

\*after C2 symmetry expansion

**Supplementary Table 2.**
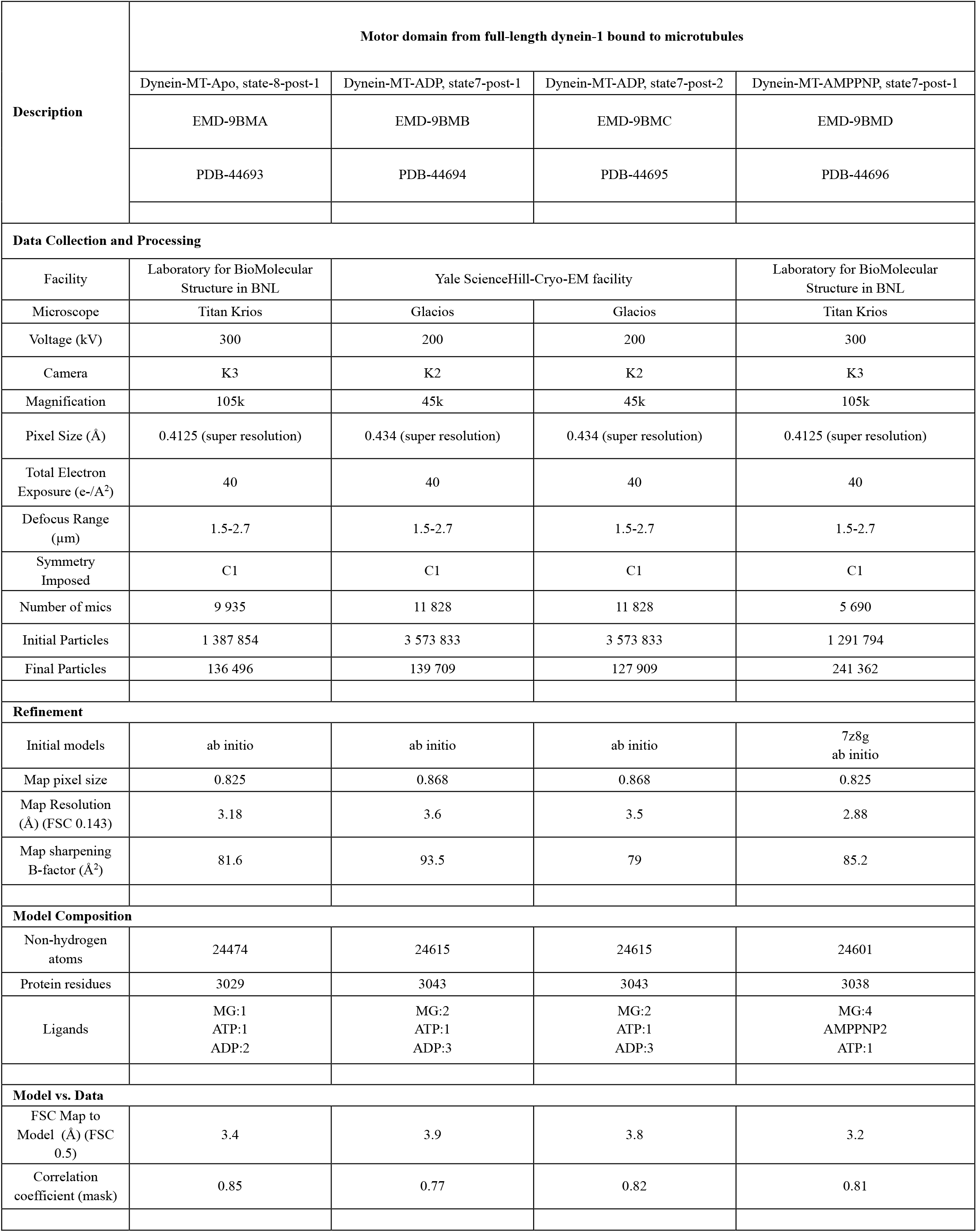
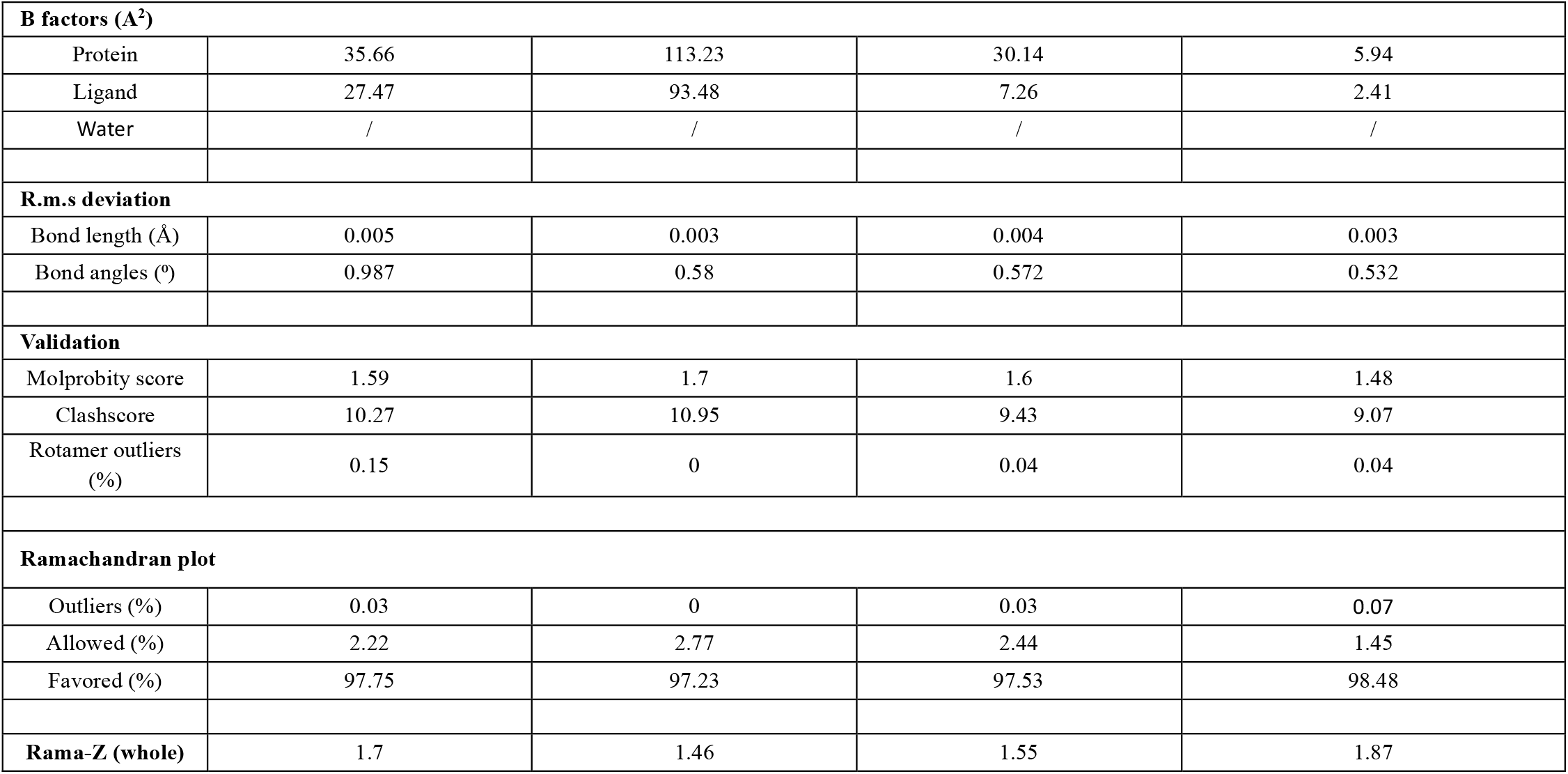
Cryo-EM data collection, refinement, and validation statistics of full-length dynein-1 bound to MTs.

**Supplementary Table 3.** Cryo-EM data collection, refinement, and validation statistics of full-length dynein-1 in the presence of 5 mM ATP-Vi with 5 mM Mg^2+^.

| Description | Full-length dynein-1 in the presence of 5 mM ATP-Vi with 5 mM Mg<sup>2+</sup> |  |  |
| --- | --- | --- | --- |
|  | State-1<br>phi-motor | State-2 | State-4 |
|  | EMD-44697 | EMD-44698 | EMD-44699 |
|  | PDB-9BMF | PDB-9BMG | PDB-9BMH |
| **Data Collection and Processing** |  |  |  |
| Facility | Yale ScienceHill-Cryo-EM facility |  |  |
| Microscope | Glacios |  |  |
| Voltage (kV) | 200 |  |  |
| Camera | K3 |  |  |
| Magnification | 45k |  |  |
| Pixel Size (Å) | 0.432 (super resolution) |  |  |
| Total Electron Exposure (e-/Å<sup>2</sup>) | 40 |  |  |
| Defocus Range (µm) | 1.5-2.7 |  |  |
| Symmetry Imposed | C1* | C1 | C1 |
| Num of mics | 6016 |  |  |
| Initial Particles | 2 554 074 |  |  |
| Final Particles | 266 214 | 69 963 | 248 329 |
| **Refinement** |  |  |  |
| Initial models |  |  |  |
| Map pixel size | 1.157 | 1.157 | 1.157 |
| Map Resolution (Å) (FSC 0.143) | 2.78 | 3.04 | 2.81 |
| Map sharpening B-factor (Å<sup>2</sup>) | 72.6 | 59.5 | 73.5 |
| **Model Composition** |  |  |  |
| Non-hydrogen atoms | 23714 | 23714 | 23,046 |
| Protein residues | 2937 | 2937 | 2,859 |
| Ligands | AOV:1<br>MG:2<br>ATP:1<br>ADP:2 | AOV:1<br>MG:2<br>ATP:1<br>ADP:2 | AOV:1<br>MG:2<br>ATP:1<br>ADP:2 |
| **Model vs. Data** |  |  |  |
| FSC Map to Model (Å) (FSC 0.5) | 3 | 3.3 | 3 |
| Correlation coefficient (mask) | 0.84 | 0.86 | 0.84 |
| **B factors (Å²)** |  |  |  |
| Protein | 61.21 | 74.39 | 57.9 |
| Ligand | 45.53 | 55.35 | 46.15 |
| Water | / | / | / |
| **R.m.s deviation** |  |  |  |
| Bond length (Å) | 0.002 | 0.002 | 0.003 |
| Bond angles (°) | 0.462 | 0.482 | 0.52 |
| **Validation** |  |  |  |
| Molprobability score | 1.26 | 1.33 | 1.36 |
| Clashscore | 4.93 | 5.95 | 6.55 |
| Rotamer outliers (%) | 0 | 0 | 0 |
| **Ramachandran plot** |  |  |  |
| Outliers (%) | 0 | 0 | 0 |
| Allowed (%) | 1.37 | 1.43 | 1.51 |
| Favored (%) | 98.63 | 98.57 | 98.49 |
| **Rama-Z (whole)** | 1.79 | 1.54 | 1.95 |

**Supplementary Table 4.**
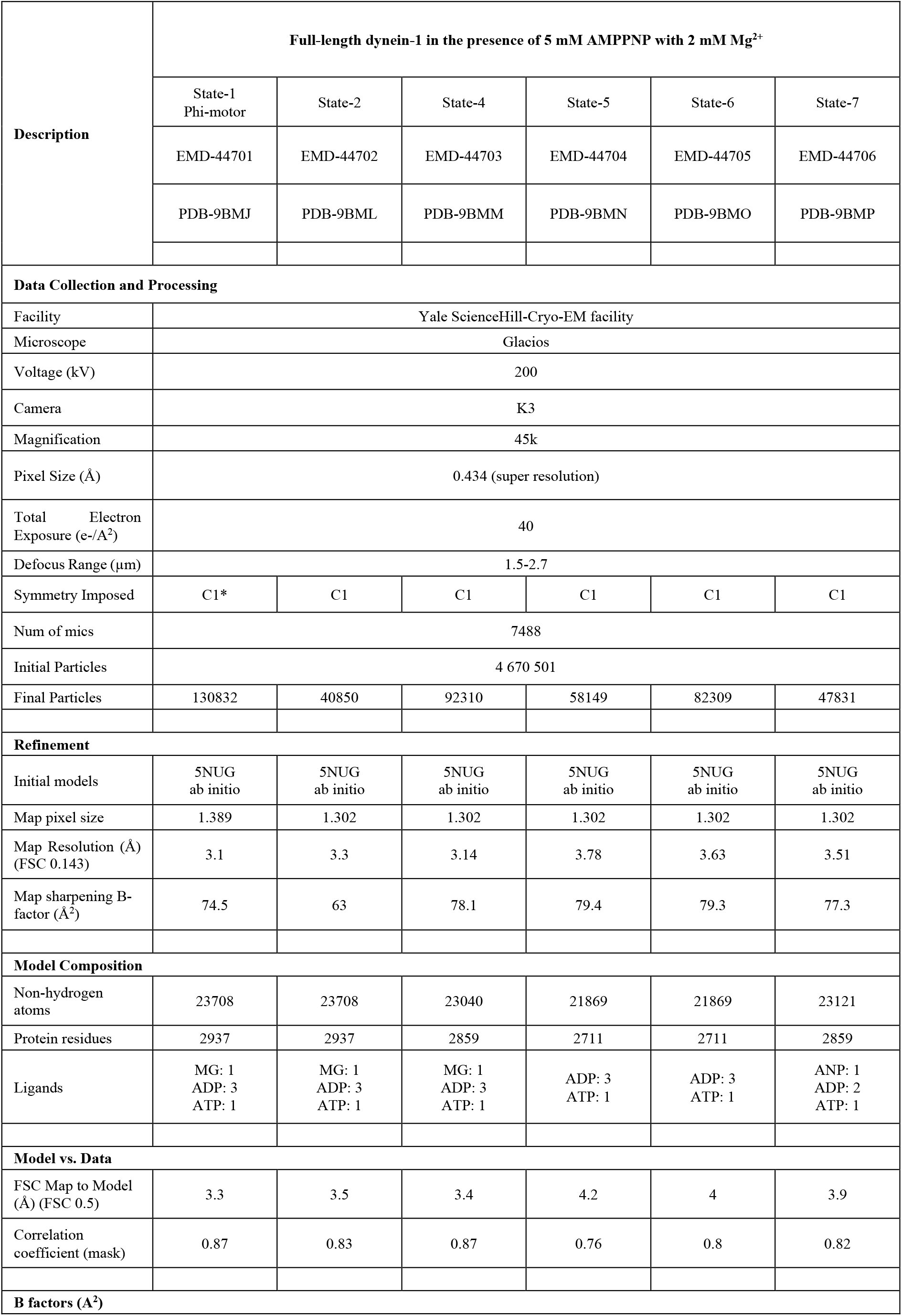
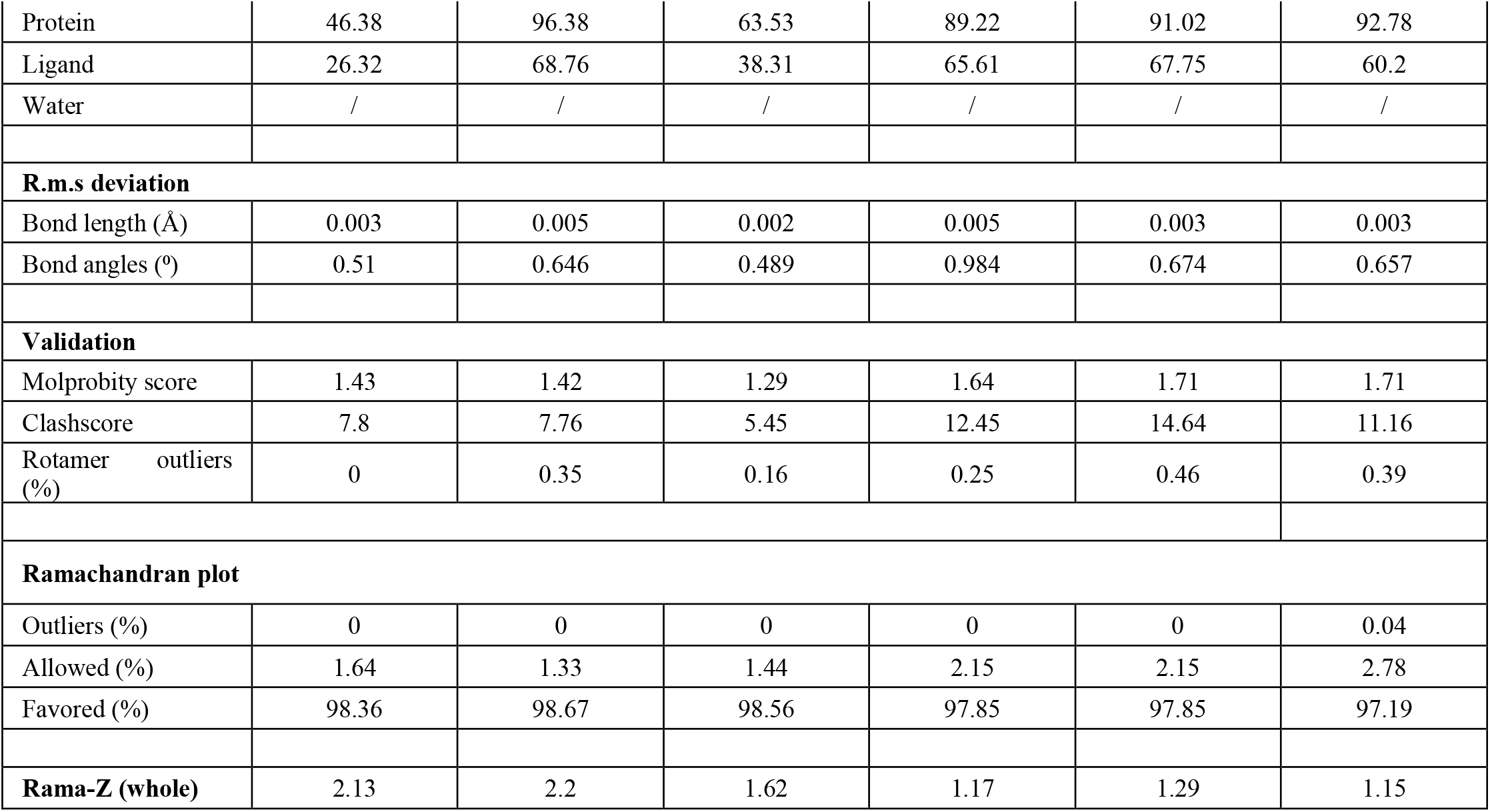
Cryo-EM data collection, refinement, and validation statistics of full-length dynein-1 in the presence of 5 mM AMPPNP with 2 mM Mg^2+^.

| Description | Full-length dynein-1 in the presence of 5 mM AMPPNP with 2 mM Mg<sup>2+</sup> |  |  |  |  |  |
| --- | --- | --- | --- | --- | --- | --- |
|  | State-1<br>Phi-motor | State-2 | State-4 | State-5 | State-6 | State-7 |
|  | EMD-44701 | EMD-44702 | EMD-44703 | EMD-44704 | EMD-44705 | EMD-44706 |
|  | PDB-9BMJ | PDB-9BML | PDB-9BMM | PDB-9BMN | PDB-9BMO | PDB-9BMP |
| Data Collection and Processing |  |  |  |  |  |  |
| Facility | Yale ScienceHill-Cryo-EM facility |  |  |  |  |  |
| Microscope | Glacios |  |  |  |  |  |
| Voltage (kV) | 200 |  |  |  |  |  |
| Camera | K3 |  |  |  |  |  |
| Magnification | 45k |  |  |  |  |  |
| Pixel Size (Å) | 0.434 (super resolution) |  |  |  |  |  |
| Total Electron Exposure (e-/Å<sup>2</sup>) | 40 |  |  |  |  |  |
| Defocus Range (µm) | 1.5-2.7 |  |  |  |  |  |
| Symmetry Imposed | C1* | C1 | C1 | C1 | C1 | C1 |
| Num of mics | 7488 |  |  |  |  |  |
| Initial Particles | 4 670 501 |  |  |  |  |  |
| Final Particles | 130832 | 40850 | 92310 | 58149 | 82309 | 47831 |
| Refinement |  |  |  |  |  |  |
| Initial models | 5NUG<br>ab initio | 5NUG<br>ab initio | 5NUG<br>ab initio | 5NUG<br>ab initio | 5NUG<br>ab initio | 5NUG<br>ab initio |
| Map pixel size | 1.389 | 1.302 | 1.302 | 1.302 | 1.302 | 1.302 |
| Map Resolution (Å)<br>(FSC 0.143) | 3.1 | 3.3 | 3.14 | 3.78 | 3.63 | 3.51 |
| Map sharpening B-factor (Å<sup>2</sup>) | 74.5 | 63 | 78.1 | 79.4 | 79.3 | 77.3 |
| Model Composition |  |  |  |  |  |  |
| Non-hydrogen atoms | 23708 | 23708 | 23040 | 21869 | 21869 | 23121 |
| Protein residues | 2937 | 2937 | 2859 | 2711 | 2711 | 2859 |
| Ligands | MG: 1<br>ADP: 3<br>ATP: 1 | MG: 1<br>ADP: 3<br>ATP: 1 | MG: 1<br>ADP: 3<br>ATP: 1 | ADP: 3<br>ATP: 1 | ADP: 3<br>ATP: 1 | ANP: 1<br>ADP: 2<br>ATP: 1 |
| Model vs. Data |  |  |  |  |  |  |
| FSC Map to Model (Å) (FSC 0.5) | 3.3 | 3.5 | 3.4 | 4.2 | 4 | 3.9 |
| Correlation coefficient (mask) | 0.87 | 0.83 | 0.87 | 0.76 | 0.8 | 0.82 |
| B factors (Å<sup>2</sup>) |  |  |  |  |  |  |
| Protein | 46.38 | 96.38 | 63.53 | 89.22 | 91.02 | 92.78 |
| Ligand | 26.32 | 68.76 | 38.31 | 65.61 | 67.75 | 60.2 |
| Water | / | / | / | / | / | / |
| **R.m.s deviation** |  |  |  |  |  |  |
| Bond length (Å) | 0.003 | 0.005 | 0.002 | 0.005 | 0.003 | 0.003 |
| Bond angles (°) | 0.51 | 0.646 | 0.489 | 0.984 | 0.674 | 0.657 |
| **Validation** |  |  |  |  |  |  |
| Molprobrity score | 1.43 | 1.42 | 1.29 | 1.64 | 1.71 | 1.71 |
| Clashscore | 7.8 | 7.76 | 5.45 | 12.45 | 14.64 | 11.16 |
| Rotamer outliers (%) | 0 | 0.35 | 0.16 | 0.25 | 0.46 | 0.39 |
| **Ramachandran plot** |  |  |  |  |  |  |
| Outliers (%) | 0 | 0 | 0 | 0 | 0 | 0.04 |
| Allowed (%) | 1.64 | 1.33 | 1.44 | 2.15 | 2.15 | 2.78 |
| Favored (%) | 98.36 | 98.67 | 98.56 | 97.85 | 97.85 | 97.19 |
| **Rama-Z (whole)** | 2.13 | 2.2 | 1.62 | 1.17 | 1.29 | 1.15 |

**Supplementary Table 5.** Cryo-EM data collection, refinement, and validation statistics of full-length dynein-1 in the presence of 5 mM AMPPNP with 5 mM Mg^2+^.

|  |  |  |  |  |  |  |
| --- | --- | --- | --- | --- | --- | --- |
| Description | Full-length dynein-1 in the presence of 5 mM AMPPNP with 5 mM Mg<sup>2+</sup> |  |  |  |  |  |
|  | State-1<br>Phi-motor | State-2 | State-4 | State-5 | State-7-Post-2 | State-7-Post-1 |
|  | EMD-46856 | EMD-46857 | EMD-46858 | EMD-46859 | EMD-46860 | EMD-46861 |
|  | PDB-9DH5 | PDB-9DH6 | PDB-9DH7 | PDB-9DH8 | PDB-9DH9 | PDB-9DHA |
| Data Collection and Processing |  |  |  |  |  |  |
| Facility | Yale ScienceHill-Cryo-EM facility |  |  |  |  |  |
| Microscope | Glacios |  |  |  |  |  |
| Voltage (kV) | 200 |  |  |  |  |  |
| Camera | K3 |  |  |  |  |  |
| Magnification | 45k |  |  |  |  |  |
| Pixel Size (Å) | 0.434 (super resolution) |  |  |  |  |  |
| Total Electron Exposure (e-/Å<sup>2</sup>) | 40 |  |  |  |  |  |
| Defocus Range (µm) | 1.5-2.7 |  |  |  |  |  |
| Symmetry Imposed | C1* | C1 | C1 | C1 | C1 | C1 |
| Num of mics | 7488 |  |  |  |  |  |
| Initial Particles | 4 670 501 |  |  |  |  |  |
| Final Particles | 301002 | 47152 | 20077 | 116933 | 87075 | 40983 |
| Refinement |  |  |  |  |  |  |
| Initial models | 5NUG<br>ab initio | 5NUG<br>ab initio | 5NUG<br>ab initio | 5NUG<br>ab initio | 5NUG<br>ab initio | 5NUG<br>ab initio |
| Map pixel size | 1.157 | 1.157 | 1.157 | 1.157 | 1.157 | 1.157 |
| Map Resolution (Å)<br>(FSC 0.143) | 2.6 | 3.4 | 3.4 | 3.5 | 3.4 | 3.6 |
| Map sharpening B-factor (Å<sup>2</sup>) | 81 | 52 | 42 | 72 | 71 | 56 |
| Model Composition |  |  |  |  |  |  |
| Non-hydrogen atoms | 23709 | 23709 | 23040 | 21869 | 23118 | 24592 |
| Protein residues | 2937 | 2937 | 2859 | 2711 | 2858 | 3038 |
| Ligands | MG: 2<br>ADP: 3<br>ATP: 1 | MG: 2<br>ADP: 3<br>ATP: 1 | MG: 1<br>ADP: 3<br>ATP: 1 | ADP: 3<br>ATP: 1 | MG: 3<br>ANP:1<br>ADP: 2<br>ATP: 1 | MG: 1<br>ANP:1<br>ADP: 2<br>ATP: 1 |
| Model vs. Data |  |  |  |  |  |  |
| FSC Map to Model (Å) (FSC 0.5) | 2.8 | 3.7 | 3.8 | 3.8 | 3.5 | 4.0 |
| Correlation coefficient (mask) | 0.84 | 0.82 | 0.79 | 0.78 | 0.83 | 0.85 |
| B factors (Å<sup>2</sup>) |  |  |  |  |  |  |
| Protein | 11.1 | 84.69 | 33.75 | 82.62 | 55.34 | 107.86 |
| Ligand | 1.53 | 54.27 | 10.22 | 55.42 | 37.86 | 80.82 |
| Water | / | / | / | / | / | / |
| **R.m.s deviation** |  |  |  |  |  |  |
| Bond length (Å) | 0.005 | 0.003 | 0.005 | 0.004 | 0.005 | 0.003 |
| Bond angles (°) | 0.977 | 0.527 | 0.984 | 0.656 | 0.811 | 0.600 |
| **Validation** |  |  |  |  |  |  |
| Molprobability score | 1.36 | 1.43 | 1.39 | 1.71 | 1.87 | 1.72 |
| Clashscore | 6.54 | 7.85 | 6.99 | 11.26 | 14.43 | 11.08 |
| Rotamer outliers (%) | 0 | 0 | 0.03 | 0 | 0.24 | 0 |
| **Ramachandran plot** |  |  |  |  |  |  |
| Outliers (%) | 0.07 | 0 | 0 | 0 | 0.04 | 0.04 |
| Allowed (%) | 1.54 | 1.5 | 1.44 | 2.74 | 3.23 | 2.91 |
| Favored (%) | 98.40 | 98.5 | 98.56 | 97.26 | 96.73 | 97.09 |
| **Rama-Z (whole)** | 1.58 | 2.03 | 1.56 | 1.36 | 0.80 | 1.18 |

**Supplementary Table 6.**
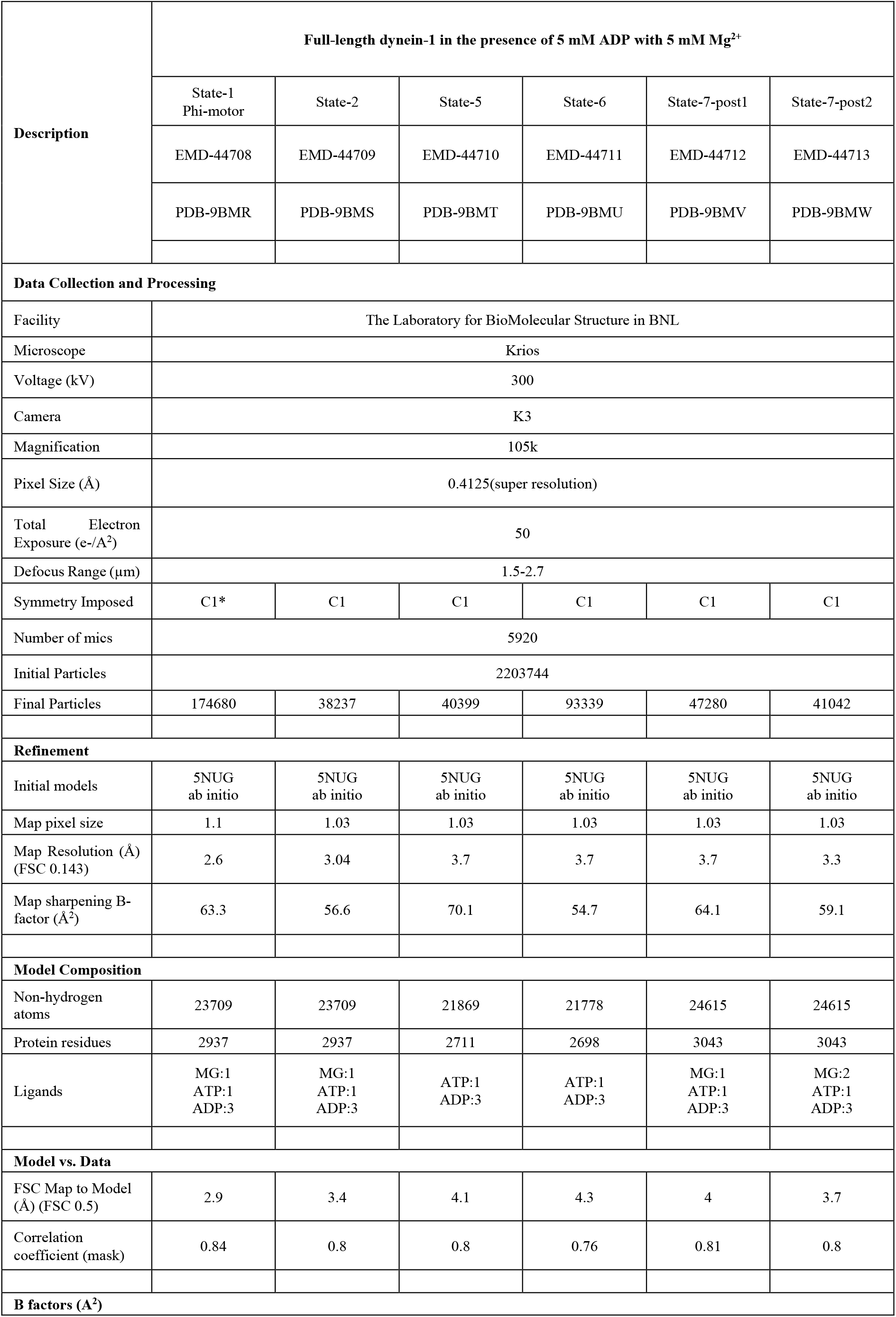
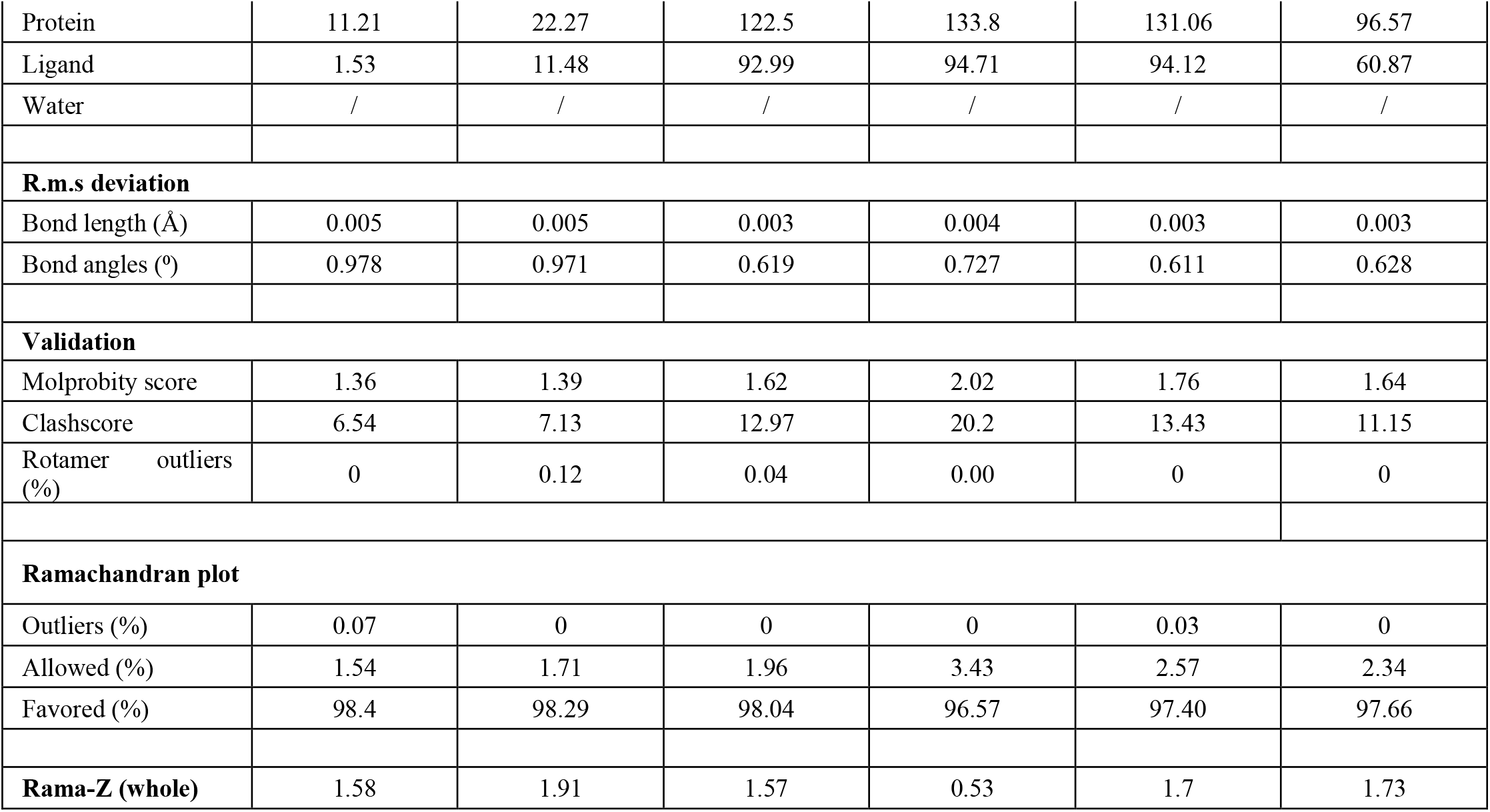
Cryo-EM data collection, refinement, and validation statistics of full-length dynein-1 in the presence of 5 mM ADP with 5 mM Mg^2+^.

**Supplementary Table 7.** Cryo-EM data collection, refinement, and validation statistics of full-length dynein-1 in apo condition with 2 mM Mg^2+^.

| Description | Full-length dynein-1 in apo buffer with 2 mM Mg<sup>2+</sup> |  |  |  |
| --- | --- | --- | --- | --- |
|  | State-1<br>Phi-motor | State-2 | State-7 | State-8 |
|  | EMD-9BMY | EMD-9BMZ | EMD-9BN0 | EMD-9BN1 |
|  | PDB-44715 | PDB-44716 | PDB-44717 | PDB-44718 |
| **Data Collection and Processing** |  |  |  |  |
| Facility | Yale ScienceHill-Cryo-EM facility |  |  |  |
| Microscope | Glacios |  |  |  |
| Voltage (kV) | 200 |  |  |  |
| Camera | K3 |  |  |  |
| Magnification | 45k |  |  |  |
| Pixel Size (Å) | 0.434 (superresolution) |  |  |  |
| Total Electron Exposure (e-/Å<sup>2</sup>) | 43 |  |  |  |
| Defocus Range (µm) | 1.5-2.7 |  |  |  |
| Symmetry Imposed | C1* | C1 | C1 | C1 |
| Number of mics | 5884 |  |  |  |
| Initial Particles | 3549872 |  |  |  |
| Final Particles | 136324 | 65852 | 67545 | 50945 |
| **Refinement** |  |  |  |  |
| Initial models | 5NUG<br>ab initio | 5NUG<br>ab initio | 5NUG<br>ab initio | 5NUG<br>ab initio |
| Map pixel size | 1.157 | 1.302 | 1.302 | 1.302 |
| Map Resolution (Å)<br>(FSC 0.143) | 2.86 | 2.97 | 3.6 | 3.8 |
| Map sharpening B-factor (Å<sup>2</sup>) | 73.6 | 58.8 | 76.7 | 71.4 |
| **Model Composition** |  |  |  |  |
| Non-hydrogen atoms | 23709 | 23709 | 23495 | 24474 |
| Protein residues | 2937 | 2937 | 2909 | 3029 |
| Ligands | MG:2<br>ATP:1<br>ADP:3 | MG:2<br>ATP:1<br>ADP:3 | MG:2<br>ATP:1<br>ADP:3 | MG:2<br>ATP:1<br>ADP:2 |
| **Model vs. Data** |  |  |  |  |
| FSC Map to Model<br>(Å) (FSC 0.5) | 3.2 | 3.2 | 4.4 | 4.2 |
| Correlation coefficient (mask) | 0.84 | 0.83 | 0.65 | 0.83 |
| **B factors (Å<sup>2</sup>)** |  |  |  |  |
| Protein | 11.21 | 22.27 | 37.72 | 141.21 |
| Ligand | 1.53 | 11.48 | 29.57 | 94.14 |
| Water | / | / | / | / |
| **R.m.s deviation** |  |  |  |  |
| Bond length (Å) | 0.005 | 0.005 | 0.006 | 0.003 |
| Bond angles (°) | 0.977 | 0.971 | 1.013 | 0.745 |
| **Validation** |  |  |  |  |
| Molprobability score | 1.37 | 1.4 | 1.66 | 1.8 |
| Clashscore | 6.64 | 7.21 | 11.55 | 16.73 |
| Rotamer outliers (%) | 0 | 0 | 0.43 | 0.41 |
| **Ramachandran plot** |  |  |  |  |
| Outliers (%) | 0.07 | 0 | 0.1 | 0 |
| Allowed (%) | 1.54 | 1.71 | 2.28 | 2.35 |
| Favored (%) | 98.4 | 98.29 | 97.62 | 97.65 |
| **Rama-Z (whole)** | 1.58 | 1.91 | 1.57 | 1.39 |

**Supplementary Table 8.** Cryo-EM data collection, refinement, and validation statistics of engineered human dynein motor domain locked in α registry.

| Description | Engineered dynein motor domain locked in $\alpha$ registry | | | |
| --- | --- | --- | --- | --- |
|  | in ATP-Vi condition |  | in ATP condition |  |
|  | Class1<br>ADP-AAA1, ADP-AAA3 | Class2<br>ADP-AAA1, ATP-AAA3 | Class1<br>ADP-AAA1, ADP-AAA3 | Class2<br>ADP-AAA1, ATP-AAA3 |
|  | EMD-44722 | EMD-44723 | EMD-44720 | EMD-44721 |
|  | PDB-9BN5 | PDB-9BN6 | PDB-9BN3 | PDB-9BN4 |
| **Data Collection and Processing** |  |  |  |  |
| Facility | Yale ScienceHill-Cryo-EM facility |  | Laboratory for BioMolecular Structure in BNL |  |
| Microscope | Glacios |  | Titan Krios |  |
| Voltage (kV) | 200 |  | 300 |  |
| Camera | K2 |  | K3 |  |
| Magnification | 36k |  | 105k |  |
| Pixel Size (Å) | 0.57 (super resolution) |  | 0.4125 (super resolution) |  |
| Total Electron Exposure (e-/Å<sup>2</sup>) | 40 |  | 40 |  |
| Defocus Range (μm) | 1.5-2.7 |  | 1.5-2.7 |  |
| Symmetry Imposed | C1 |  | C1 |  |
| Number of mics | 2466 |  | 9 498 |  |
| Initial Particles | 638 774 |  | 2 954 198 |  |
| Final Particles | 112 474 | 72 335 | 433 112 | 91 718 |
| **Refinement** |  |  |  |  |
| Initial models | ab initio | ab initio | ab initio | ab initio |
| Map pixel size | 1.143 | 1.143 | 1.146 | 1.146 |
| Map Resolution (Å) (FSC 0.143) | 3.3 | 3.4 | 2.8 | 2.9 |
| Map sharpening B-factor (Å<sup>2</sup>) | 61.4 | 53.6 | 79 | 64.4 |
| **Model Composition** |  |  |  |  |
| Non-hydrogen atoms | 23158 | 23084 | 23158 | 23084 |
| Protein residues | 2866 | 2855 | 2866 | 2855 |
| Ligands | ATP:1<br>ADP:3 | ATP:2<br>ADP:2 | ATP:1<br>ADP:3 | ATP:2<br>ADP:2 |
| **Model vs. Data** |  |  |  |  |
| FSC Map to Model (Å) (FSC 0.5) | 3.7 | 3.9 | 3.2 | 3.3 |
| Correlation coefficient (mask) | 0.8 | 0.8 | 0.75 | 0.79 |
| **B factors (Å<sup>2</sup>)** |  |  |  |  |
| Protein | 82.16 | 119.44 | 19.63 | 60.63 |
| Ligand | 56.75 | 91.8 | 5.58 | 33.32 |
| Water | / | / | / | / |
| **R.m.s deviation** |  |  |  |  |
| Bond length (Å) | 0.002 | 0.003 | 0.004 | 0.004 |
| Bond angles (°) | 0.584 | 0.607 | 0.672 | 0.709 |
| **Validation** |  |  |  |  |
| Molprobity score | 1.69 | 1.72 | 1.8 | 1.71 |
| Clashscore | 8.89 | 11.09 | 11.03 | 11.5 |
| Rotamer outliers (%) | 0.04 | 0 | 0.04 | 0.04 |
| **Ramachandran plot** |  |  |  |  |
| Outliers (%) | 0.04 | 0.14 | 0.04 | 0.14 |
| Allowed (%) | 3.29 | 2.75 | 3.57 | 2.57 |
| Favored (%) | 96.67 | 97.11 | 96.39 | 97.29 |
| **Rama-Z (whole)** | 0.82 | 1.06 | 0.75 | 0.81 |

